# The architecture of the centriole cartwheel-containing region revealed by cryo-electron tomography

**DOI:** 10.1101/2020.07.21.068882

**Authors:** Nikolai Klena, Maeva Le Guennec, Anne-Marie Tassin, Hugo van den Hoek, Philipp S. Erdmann, Miroslava Schaffer, Stefan Geimer, Gabriel Aeschlimann, Lubomir Kovacik, Kenneth N. Goldie, Henning Stahlberg, Benjamin D. Engel, Virginie Hamel, Paul Guichard

## Abstract

Centrioles are evolutionarily conserved barrels of microtubule triplets that form the core of the centrosome and the base of the cilium. In the proximal region of the centriole, nine microtubule triplets attach to each other via A-C linkers and encircle a central cartwheel structure, which directs the early events of centriole assembly. While the crucial role of the proximal region in centriole biogenesis has been well documented in many species, its native architecture and evolutionary conservation remain relatively unexplored. Here, using cryo-electron tomography of centrioles from four evolutionarily distant species, including humans, we report on the architectural diversity of the centriolar proximal cartwheel-bearing region. Our work reveals that the cartwheel central hub, previously reported to have an 8.5 nm periodicity in *Trichonympha*, is constructed from a stack of paired rings with an average periodicity of ∼4 nm. In all four examined species, cartwheel inner densities are found inside the hub’s ring-pairs. In both *Paramecium* and *Chlamydomonas*, the repeating structural unit of the cartwheel has a periodicity of 25 nm and consists of three ring-pairs with 6 radial spokes emanating and merging into a single bundle that connects to the triplet microtubule via the pinhead. Finally, we identified that the cartwheel is indirectly connected to the A-C linker through a flexible triplet-base structure extending from the pinhead. Together, our work provides unprecedented evolutionary insights into the architecture of the centriole proximal region, which underlies centriole biogenesis.

## Introduction

Centrioles and basal bodies (hereafter referred to as centrioles for simplicity) are cytoskeletal organelles, typically 450 to 550 nm in length and ∼250 nm in outer diameter, which are present in most eukaryotic cells and play organizing roles in the assembly of cilia, flagella and centrosomes (Gönczy, 2012; Nigg and Raff, 2009; Winey and O’Toole, 2014). Centrioles are characterized by a near-universal nine-fold radial arrangement of triplet microtubules that contain a complete 13-protofilament A-microtubule and incomplete B- and C-microtubules, each composed of 10 protofilaments (Guichard et al., 2013). Centrioles are polarized along their proximal to distal axis, with distinct structural features along their length. The proximal region is defined by the presence of the cartwheel structure, which serves as a seed for centriole formation and is thought to impart nine-fold symmetry to the entire organelle (Gönczy, 2012; Hilbert et al., 2016; Hirono, 2014; Nakazawa et al., 2007; Strnad and Gönczy, 2008). In most of the species, the cartwheel stays within the centriole after maturation, however, it is no longer present in mature human centrioles (Azimzadeh and Bornens, 2007). The native architecture of the proximal region, and in particular of the cartwheel, was revealed by cryo-electron tomography (cryo-ET) of the *Trichonympha* centriole. Owing to its exceptionally long proximal region, many structural repeats could be sampled for subtomogram averaging, revealing the overall 3D structure of the cartwheel for the first time (Guichard et al., 2013, 2012). The *Trichonympha* cartwheel was observed to be built from a hub of stacked rings spaced every 8.5 nm. Radial spokes, emanating from two adjacent rings, merged at the pinhead near the microtubule triplet to form a repeating structural unit with a periodicity of 17 nm. Moreover, this study demonstrated that each *Trichonympha* hub ring could accommodate nine homodimers of SAS-6, a protein that is essential for cartwheel assembly across eukaryotes (Kitagawa et al., 2011; van Breugel et al., 2014, 2011). Unexpectedly, a CID, for Cartwheel Inner Densities, was also identified at the center of the hub ring. This CID contacts the hub ring at nine locations and has been hypothesized to be *Trichonympha*-specific, as it has never been observed in other species, possibly due to lack of resolution. In this respect, the CID has been proposed to facilitate TaSAS-6 oligomerization or confer additional mechanical stability to these exceptional long centrioles, which are subjected to strong forces inside the intestine of the host termite (Guichard et al., 2018, 2013).

In the proximal region, the cartwheel is connected to the pinhead, which bridges the cartwheel to the A-microtubule of the microtubule triplet (Dippell, 1968; Hirono, 2014). This connection is thought to be partially composed of Bld10p/Cep135 proteins, which can interact both with SAS-6 and tubulin (Carvalho-Santos et al., 2012; Guichard et al., 2017; Hiraki et al., 2007a; Kraatz et al., 2016). In addition to the cartwheel/pinhead ensemble, adjacent microtubule triplets in the proximal region are also connected by the A-C linker. Cryo-ET combined with subtomogram averaging has revealed distinct structures of the A-C linker in *Trichonympha* and *Chlamydomonas reinhardtii* (Guichard et al., 2013; Li et al., 2019). In *Trichonympha*, the structure consists of the A-link, which is laterally inclined and contacts the A-tubule at the A8 protofilament, and the C-link, which connects to the C-tubule at the C9 protofilament. Overall, the *Trichonympha* A-C linker displays a longitudinal periodicity of 8.5 nm. In contrast, the A-C linker in *C. reinhardtii* is a crisscross-shaped structure composed of a central trunk region from which two arms and two legs extend to contact the A- and C-tubules (Li et al., 2019). Whereas these two studies provide major advances in our understanding of A-C linker organization, they also clearly highlight structural divergence between *Trichonympha* and *C. reinhardtii* centrioles.

The question thus arises as to the evolutionary conservation of the centriole’s proximal region, including characteristic structures such as the A-C linker and the cartwheel’s hub, CID and radial spokes. In particular, the structure of the cartwheel remains unexplored beyond *Trichonympha*. A more universal description of the proximal region is important for understanding of how these structures direct centriole biogenesis. Here, we use cryo-ET to tackle this fundamental question using four evolutionary distant species: *Chlamydomonas reinhardtii, Paramecium tetraurelia, Naegleria gruberi*, and humans.

## Results

### *In situ* structural features of the cartwheel in *Chlamydomonas* centrioles

The power of biodiversity proved extremely useful for resolving the first 3D architecture of the cartwheel within the exceptionally long proximal region of *Trichonympha* centrioles (Guichard et al., 2012). This study identified the CID as well as an 8.5 nm longitudinal periodicity along the central hub of the cartwheel. Whether these structural features hold true in other species is an open question that we address here by analyzing the cartwheel of the green algae *C. reinhardtii*, a canonical model for centriole biology with similar centriole structure and protein composition to humans (Hamel et al., 2017; Keller et al., 2005; Keller and Marshall, 2008; Li et al., 2011). However, extracting centrioles from cells can limit the analysis of these fragile structures, as exemplified by the loss of the cartwheel during a study of isolated *C. reinhardtii* centrioles (Li et al., 2011). In addition, the >300 nm thick vitreous ice surrounding uncompressed centrioles on an EM grid reduces the signal and contrast of cryo-ET (Kudryashev et al., 2012), making it difficult to resolve fine details in the relatively small cartwheel structure (Guichard et al., 2018). We therefore decided to analyze the *C. reinhardtii* cartwheel *in situ* using a cryo-focused ion beam (cryo-FIB) milling approach, which creates thin 100-150 nm sections of the native cellular environment in a vitreous state (Schaffer et al., 2017). Combining this approach with new direct electron detector cameras (Grigorieff, 2013), it was possible for us to visualize the centriole and cartwheel with unprecedented clarity and structural preservation.

As shown in Figure 1A-B, *in situ* cryo-ET clearly revealed both mature centrioles and procentrioles, providing the first observation of the centriole’s cartwheel-bearing region in its native environment. The cartwheel’s structural features were analyzed in both types of centrioles (Figures 1C-H, S1 and S2). Strikingly, we found that the cartwheel’s central hub has an average longitudinal periodicity of 4.0 nm in both mature centrioles and procentrioles, distinct from the 8.5 nm periodicity originally described in *Trichonympha* (Guichard et al., 2012) (Figures 1H and S1A, D, G). Moreover, we noticed pronounced densities inside the central hub that were reminiscent of the CID originally described in *Trichonympha*, suggesting that this structure is not *Trichonympha*-specific but rather is a conserved feature of the cartwheel (Figure 1). Several CIDs in *C. reinhardtii* are spaced along the lumen of the central hub, forming an 8.7 nm periodicity on average, in mature centrioles and procentrioles (Figures 1H and S1B, E), similar to *Trichonympha*.

**Figure 1.**
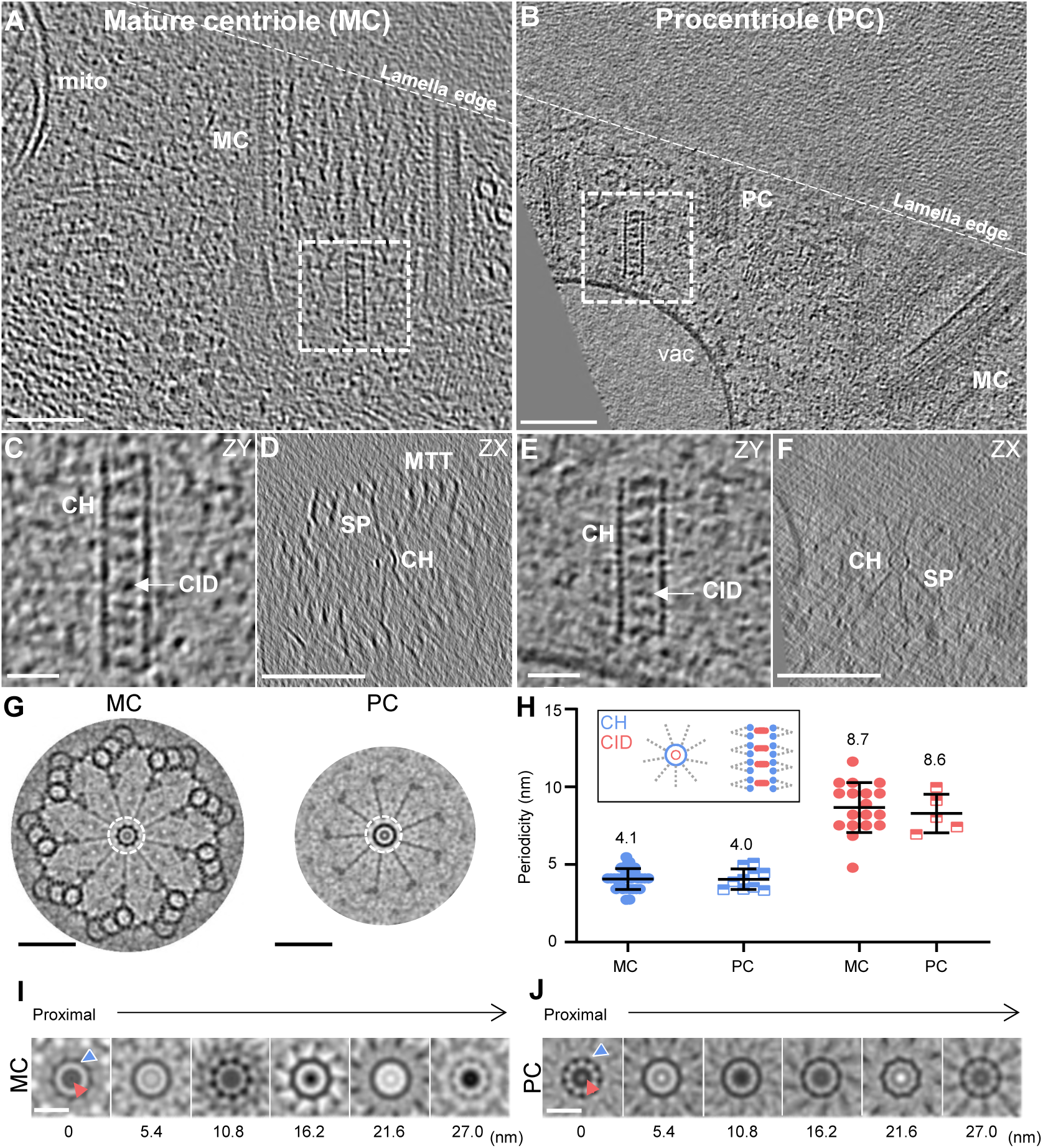
*In situ* cryo-ET reveals the native cartwheel structure in *C. reinhardtii* centrioles. (A, B) *In situ* cryo-electron tomogram displaying the proximal region of a mature mother centriole (A) and procentriole (B). Mature centriole, MC; procentriole, PC; mitochondria, mito; vacuole, vac; white dashed line, lamella edge. Scale bars, 100 nm. (C, E) Side view z-projection of cartwheels containing the central hub and several CIDs from a mature centriole (C) and a procentriole (E). Central hub, CH; cartwheel inner densities, CID. Scale bars, 20 nm. (D, F) Cross sections of the cartwheel-containing regions from a mature centriole (D) and a procentriole (F). Microtubule triplet, MTT; spokes, SP. Scale bars, 200 nm. (G) Nine-fold symmetrized cross sections of the cartwheel-containing region from a mature centriole (left side) and a procentriole (right side). Dashed white circle, central hub. Scale bars, 100 nm. (H) Longitudinal periodicity measurements of the central hub and CIDs. Central hub, blue; CID, red. Mean values are displayed above the data range. (I, J) Nine-fold symmetrized central hub z-projections, starting at the proximal end of the cartwheel and continuing distally along the cartwheel by 5.4 nm steps in a mature centriole (I) and a procentriole (J). Red arrow, CID; blue arrow; central hub.

To investigate whether the discrepancy we observed in central hub periodicity was accompanied by other differences in cartwheel structure, we measured features of the cartwheel such as the central hub diameter as well as the distances from the hub to D1 and D2, two densities previously described on the cartwheel spokes of *C. reinhardtii* centrioles (Guichard et al., 2017) in both mature centrioles and procentrioles (Figure S3A-F). Similar to previous measurements, we found that the central hub is ∼21 nm in diameter (peak-to-peak from the intensity plot profile through the hub), and the D1 and D2 densities are positioned ∼ 36 nm and ∼ 47 nm from the external edge of the cartwheel hub, respectively. These measurements suggest that only the longitudinal periodicity of the central hub differs in the *in situ C. reinhardtii* centrioles (Figure S3A-F).

While most of the cartwheel’s structural features, including the CID, are conserved between *Trichonympha* and *C. reinhardtii*, the periodicity of the central hub appears to diverge. This discrepancy poses the important question of how conserved the architecture of the cartwheel-containing region is between species. Moreover, as cartwheel periodicity was previously only measured in isolated centrioles, this raises the possibility that cartwheel periodicity may be affected during purification.

### Conservation of the cartwheel’s structural features in *Paramecium, Naegleria* and humans

To address these questions, we analyzed the proximal region of isolated centrioles from three different species. Centrioles were purified from *P. tetraurelia, N. gruberi* and human KE37 leukemia acute lymphoblastic T cells, vitreously frozen onto EM grids, and then imaged by cryo-ET (Figure 2A-I). Despite the high level of noise expected in cryo-ET of isolated centrioles and the previously observed strong compression of *N. gruberi* and human centrioles (Greenan et al., 2018; Guichard et al., 2010; Le Guennec et al., 2020) that affects cartwheel integrity, we could reliably measure the central hub periodicity in each of these species. Strikingly, we found that the longitudinal periodicity of the central hub is similar to the *C. reinhardtii in situ* cartwheel, with average periodicities of 4.3 +/− 0.38 nm, 4.4 +/− 0.53 nm and 4.2 +/− 0.68 nm in *P. tetraurelia, N. gruberi* and human, respectively (Figure 2J). Moreover, we observed that CID structures are present in every species, forming a periodicity along the central hub of 8.4 +/− 1.25 nm, 8.3 +/− 1.83 nm and 8.1 +/− 2.46 nm (Figure 2A-J and Figure S3G-O). These results indicate that structural features of the *C. reinhardtii* cartwheel seem to be conserved, including the central hub’s ∼4.2 nm periodicity, as well as the presence of CIDs every ∼8.4 nm. Moreover, these measurements demonstrate that the discrepancy between *Trichonympha* and *C. reinhardtii* is possibly not due to purification artifacts, as the other isolated centrioles also display ∼4 nm periodicities along their central hubs.

**Figure 2.**
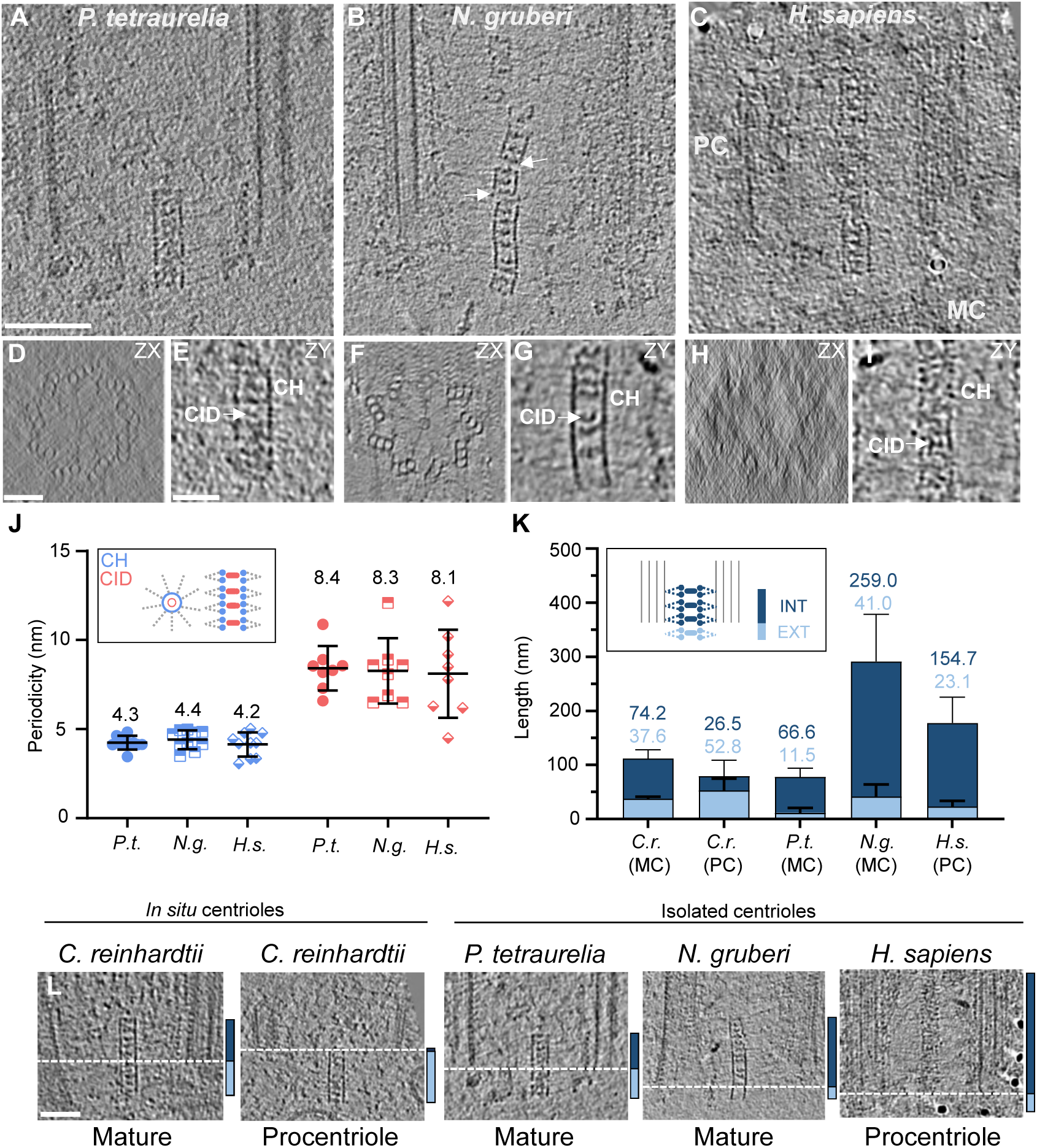
Cryo-ET of isolated centrioles from *P. tetraurelia, N. gruberi*, and *H. sapiens* reveals novel cartwheel periodicities. (A, B, C) Cryo-electron tomograms of the proximal regions of a *P. tetraurelia* centriole (A), a *N. gruberi* centriole, (B) and a *H. sapiens* procentriole (C). White arrows denote a broken cartwheel; procentriole, PC; mature centriole, MC; Scale bar, 100 nm. Note that *N. gruberi* and *H. sapiens* centrioles were heavily compressed during the cryo-EM preparation, as previously described (Guichard et al., 2010). The periodicities in *N. gruberi* and *H. sapiens* centrioles were measured only on parts that were not damaged. (D, F, H) Cross sections from cartwheel-containing regions of *P. tetraurelia* (D), *N. gruberi* (F), and *H. sapiens* (H) centrioles. Scale bar, 50 nm. (E, G, I) Zoomed side views of cartwheels from *P. tetraurelia* (E), *N. gruberi* (G), and *H. sapiens* (I), displaying the central hub (CH) and several cartwheel inner densities (CIDs), white arrow. Scale bar, 25 nm. (J) Longitudinal periodicity of the central hub and CIDs in *P. tetraurelia, N. gruberi*, and *H. sapiens*. Mean value displayed above range. (K, L) Proximal protrusion of the cartwheel beyond the microtubule triplets in *C. reinhardtii, P. tetraurelia, N. gruberi*, and *H. sapiens*. Internal cartwheel inside the microtubule barrel, dark blue (INT); external cartwheel beyond the microtubule wall, light blue (EXT). Mean values are displayed above the range (K). Start of the microtubule wall is delineated by dashed white line (L). Scale bar, 50 nm.

Interestingly, in tomograms of both *in situ* and isolated centrioles, we observed that the position of the cartwheel did not fully correlate with the position of the microtubule triplets. In all four species, the cartwheels protruded proximally 10-40 nm beyond the microtubule wall (Figures 1A, B and 2K, L). In *C. reinhardtii*, which enabled observations of assembling and mature centrioles within the same cells, the cartwheel extension was more prominent in procentrioles, with 67% of the cartwheel protruding in contrast to 34% in mature centrioles (Figure 2K). Until now, this proximal extension of the cartwheel has only been reported in isolated *C. reinhardtii* procentrioles (Geimer and Melkonian, 2004; Guichard et al., 2017). Our *in situ C. reinhardtii* tomograms demonstrate that the cartwheel extension is not an artifact of purifying centrioles, but rather occurs within the native cellular environment. We further corroborated this conclusion with serial sections of resin-embedded *N. gruberi* cells, which show the cartwheel protruding beyond the proximal end of the microtubule triplets in both assembling and mature centrioles (Fig. S4). The cartwheel extension is consistent with fluorescence microscopy localization of cartwheel components CrSAS-6 and Bld10p, which extend from the centriole’s proximal region to ∼60 nm below the proximal-most acetylated tubulin signal in mature *C. reinhardtii* centrioles (Hamel et al., 2017). Additionally, this proximal extension corroborates 3D-SIM-FRAP analysis of SAS-6-GFP in *Drosophila*, showing that the cartwheel may grow from its proximal end (Aydogan et al., 2018). Taking these data together, we conclude that the cartwheel protrusion is not a consequence of biochemical isolation but rather is an evolutionarily conserved structural feature that may relate to early events in centriole assembly.

### 3D architecture of the cartwheel in *Paramecium* and *Chlamydomonas*

Given the intriguing 4.0 nm periodicity of the central hub revealed in our study, which differs from the previously reported periodicity in *Trichonympha* (Guichard et al., 2012), we decided to take a closer look at the cartwheel architecture in both *P. tetraurelia* and *C. reinhardtii* centrioles. As explained above, resolving the cartwheel structure in these species represents a major challenge, as the cartwheel length is about 40-times shorter than the exceptionally long *Trichonympha* cartwheel, limiting the number of repeat units available for subtomogram averaging. Nevertheless, we undertook this task with a low number of subvolumes, increasing the contrast of the central hub and emanating radial spokes. From 8 *P. tetraurelia* tomograms, we performed subtomogram averaging on 235 boxes and symmetrized the obtained map. A projection of the reconstructed *P. tetraurelia* cartwheel is shown in Figure 3A, where the CID, the central hub, and its emanating radial spokes are clearly visible. Careful inspection of a longitudinal section through the averaged volume confirmed the presence of CIDs every 8.6 nm inside the central hub (Figures 2J and 3C). Intriguingly, we found that the central hub is constructed from pairs of rings (Figure 3B-C, light blue arrowheads). These ring-pairs have an inter-ring distance of 3.1 nm and stack on each other with 5.5 nm between adjacent ring-pairs, resulting in the average periodicity of ∼4.2 nm along the central hub (Figures 2J and 3C). We observed that two small densities (Figure 3D, red arrows) emanate from each ring-pair (red circles) and fuse into one radial spoke (white arrows) that in turn merges with two other fused spokes to form a single structure ∼37 nm from the central hub surface, a distance that corresponds to the D1 density (Figure 3D, blue arrow; Figure S5A, blue arrows). The three ring-pairs that share fused spokes are repeated three to four times along the cartwheel length, with a longitudinal distance of ∼25 nm between merged spokes (Figure 3B and Figure S5A, blue arrows), suggesting that this represents the repeating structural unit of the cartwheel. We also noted that the emanating spokes are slightly tilted (Figure 3D, 3J and Figure S5A, green dotted lines), possibly reflecting a twist in the molecular interaction underlying spoke fusion. Interestingly, we found that the CIDs are positioned at the center of each ring-pair (Figure 3C), suggesting that they could be important for the ring-pair’s formation or stability. Importantly, all these features can also be seen within the raw data (Figure S6A, B), indicating that they are not a result of the averaging procedure.

**Figure 3.**
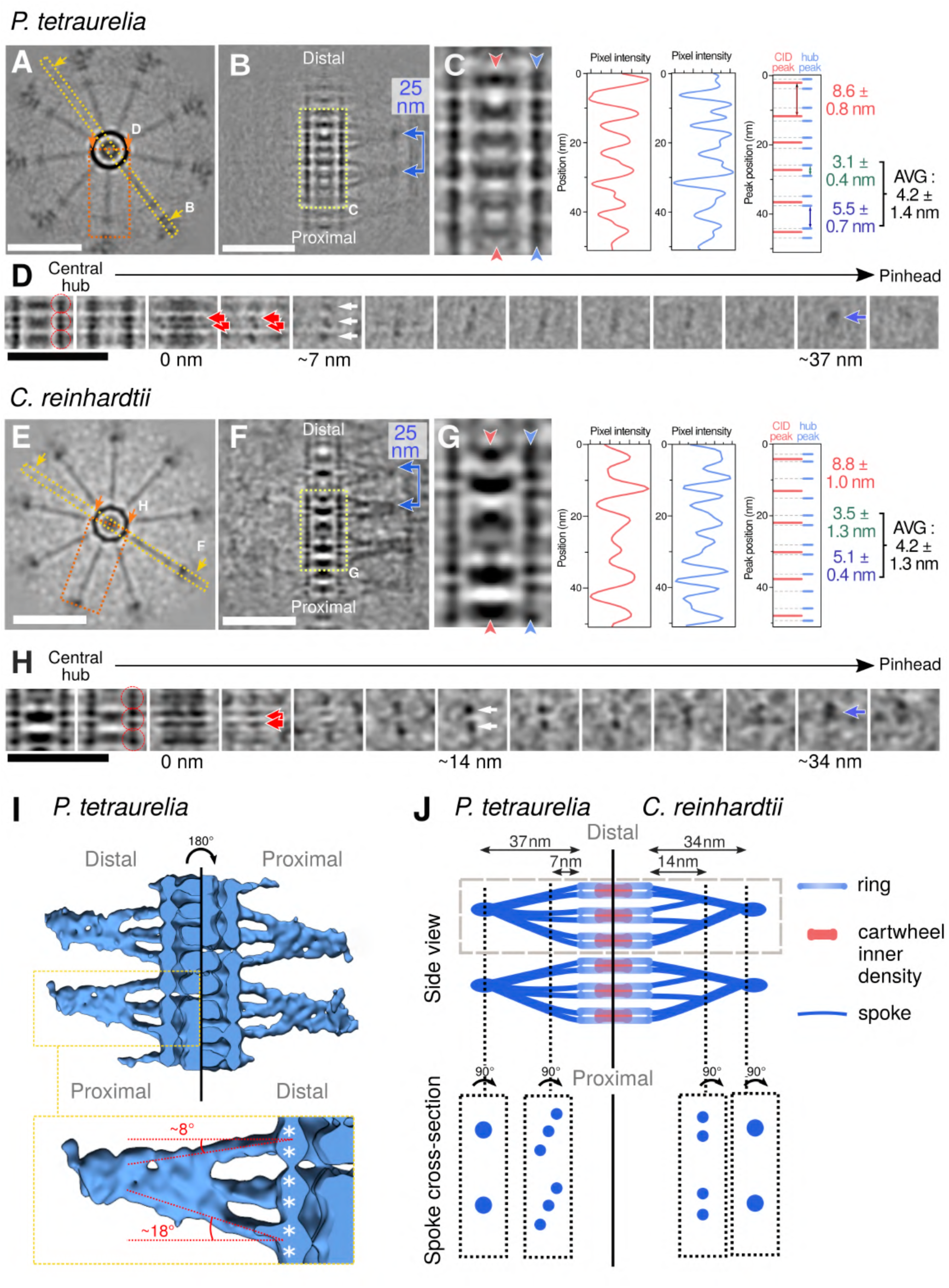
Subtomogram averaging of *P. tetraurelia* and *C. reinhardtii* cartwheels reveals novel cartwheel structural organization. (A, E) Top views of cartwheel reconstructions from *P. tetraurelia* (A) and *C. reinhardtii* (E). Scale bars, 50 nm. Dashed yellow line with arrows denotes the central hub-focused reslice shown in panels B and F; dashed red box with arrows denotes the spoke-focused reslice shown in panels D and H. (B, F) Reslice of central hub-containing region with spokes in *P. tetraurelia* (B) and *C. reinhardtii* (F). Scale bars, 50 nm. Dashed light yellow line denotes the zoomed view shown in panels C and G, blue line with arrows indicates the ∼25 nm repeat distance between merged spokes. (C, G) Zoomed view displaying periodic repeats of the central hub (CH) and several cartwheel inner densities (CIDs) in *P. tetraurelia* (C) and *C. reinhardtii* (G). CID, red arrowheads and red plot profile; CH, blue arrowheads and blue plot profile. Overlay between CIDs and CH peaks plotted on the right in red and blue. Mean distance between CIDs peaks, red +/− S.E.M. Distances between CH peaks split into two distinct populations: smaller (within a ring-pair), green +/− S.E.M; larger (between ring-pairs), blue +/− S.E.M; average periodicity; black +/− S.E.M. (D, H) Serial z-projections of ∼4 nm thickness from one cartwheel repeat unit of *P. tetraurelia* (D) and *C. reinhardtii* (H). Left-most z-projections display the central hub, right-most projection shows the microtubule wall. Red circles delineate one ring-pair. Red arrows mark individual spokes. White arrows mark merged spokes. Blue arrow indicates the final merged spoke (D1 density). Scale bar, 50 nm. (I) Three-dimensional rendering of the cartwheel reconstruction from *P. tetraurelia*. Left side, cartwheel oriented along the correct proximal-distal axis; right side, 180° inverted proximal-distal axis, showing the asymmetry of spoke inclination. Dashed yellow box, inset of one spoke unit, with the major and minor tilt angles of the spokes relative to the central hub. White asterisks denote subunits of ring-pairs. (J) Model of *P. tetraurelia* (left side) and *C. reinhardtii* (right side) cartwheel structures. Dashed grey box denotes one repeat unit of the cartwheel, dashed black lines and boxes display cross sections of spokes.

Next, we performed a similar analysis on *C. reinhardtii* mature centrioles (Figure 3E), using 102 subvolumes from 5 *in situ* tomograms and then applied symmetrization. Interestingly, we found that the cartwheel’s repeating structural unit is also composed of three ring-pairs, with 3.5 nm inter-ring spacing and 5.1 nm spacing between ring-pairs (Figure 3E-G, blue arrowhead in G), leading to the observed ∼4.2 nm periodicity along the central hub. Each repeating unit also had six emanating spokes (Figure 3H and Figure S5B red arrows); however, these spokes were organized differently than in *P. tetraurelia* cartwheels, merging into two spokes ∼14 nm from the central hub (Figure 3H and Figure S5B white arrows) and further fusing into a single unit ∼34 nm from the hub (Figure 3H and Figure S5B blue arrows). Similar to *P. tetraurelia*, the repeating unit of the central hub has a periodicity of ∼ 25 nm (Figure 3F and Figure S5B blue arrows). In *C. reinhardtii* cartwheels, CIDs are positioned 8.8 nm apart, inside ring-pairs (Figure 3G). As for *P. tetraurelia*, we confirmed that these *C. reinhardtii* features could be seen in the raw data (Figure S6C, D) and were not a result of the averaging. We also noticed in raw tomograms that some regions were devoid of CIDs, suggesting that their positioning might be stochastic (Figure S6D, white arrowhead).

Together, these results demonstrate that both species have an overall similar cartwheel organization, with some species-specific differences in the radial spokes that possibly reflect either a different modality of assembly or some divergence at the molecular level. Moreover, we also noticed that the repeating structural unit described here displays a polarity from proximal to distal that is defined by the angle of the emanating spokes, which is strikingly apparent in the *P. tetraurelia* average (Figure 3I-J).

Next, we investigated how the observed discrepancy in central hub periodicity could arise between *C. reinhardtii* / *P. tetraurelia* and *Trichonympha*. We hypothesized that the resolution improvement from using a direct electron detector might have helped reveal features that were not visible in the previous study of *Trichonympha* centrioles. To test this idea, we applied a bandpass filter to decrease the resolution of the *P. tetraurelia* subtomogram average to that of the *Trichonympha* map (38 Å) (Figure S5C, D). At this resolution, the *P. tetraurelia* ring-pairs appear to be single rings, leading to a global 8.6 nm periodicity along the central hub as originally described in *Trichonympha*. This result suggests that the *Trichonympha* cartwheel most likely also exhibits the same ∼4 nm ring-pair periodicity as *P. tetraurelia* and *C. reinhardtii*, but this could not be retrieved in earlier studies primarily due to resolution limitations of the detectors used for imaging. However, we also noticed that the spoke organization appears different between *Trichonympha* and *C. reinhardtii* / *P. tetraurelia* cartwheels, suggesting variability of molecular organization between species.

### Defining the structural features of the proximal region

We next focused on charting the overall organization present in the cartwheel-containing region of *P. tetraurelia* and *C. reinhardtii* centrioles to better understand how the cartwheel is connected to the microtubules, as well as to check whether the structural features are conserved between species (Figure 4). As subtomogram averaging might average out non-periodic structures, we first analyzed the raw tomograms by systematically extracting cross sections of centrioles from both species at different positions along the proximal-to-distal axis and then applying nine-fold symmetrization to improve the contrast using centrioleJ (Guichard et al., 2013) (Figure 4B, H). Starting from the proximal side, several previously described structural features could be resolved, including the cartwheel (blue arrow), the pinhead (magenta arrow), the A-C linker (turquoise arrow) and the beginning of the inner scaffold (orange arrow) that defines the central core region of the centriole (Figure 4B, C and H, I, F). We also noticed a linker between the pinhead structure and the A-C linker (Figure 4C panels (III, IV) and 4I panels (III, IV), light green arrow). This linker is reminiscent of the triplet base structure originally described in human, mouse, and Chinese hamster centrioles (Vorobjev and Chentsov, 1980) and also detected in *Trichonympha* centrioles (Gibbons and Grimstone, 1960). We therefore conclude that the triplet base is an evolutionarily conserved structural feature of the centriole’s cartwheel-bearing region. Interestingly, in contrast to the A-C linker (Figure 4C, panel (VI) and 4I, panel (VI)), the pinhead structure does not co-exist with the inner scaffold, suggesting that the later replaces the former (Figure 4D, E, J-N and Figure S7 A, D). We also noticed in the most distal part of the proximal region that the pinhead structure is present without the cartwheel in *P. tetraurelia* centrioles (Figure 4B, C panel (IV), D, E, and Figure S7A, S7D). Finally, we observed in the two *in situ C. reinhardtii* procentrioles that the A-C linker covers the entire length of the growing microtubule triplets, while the pinhead and cartwheel seem to display variable lengths (Figure S7G).

**Figure 4.**
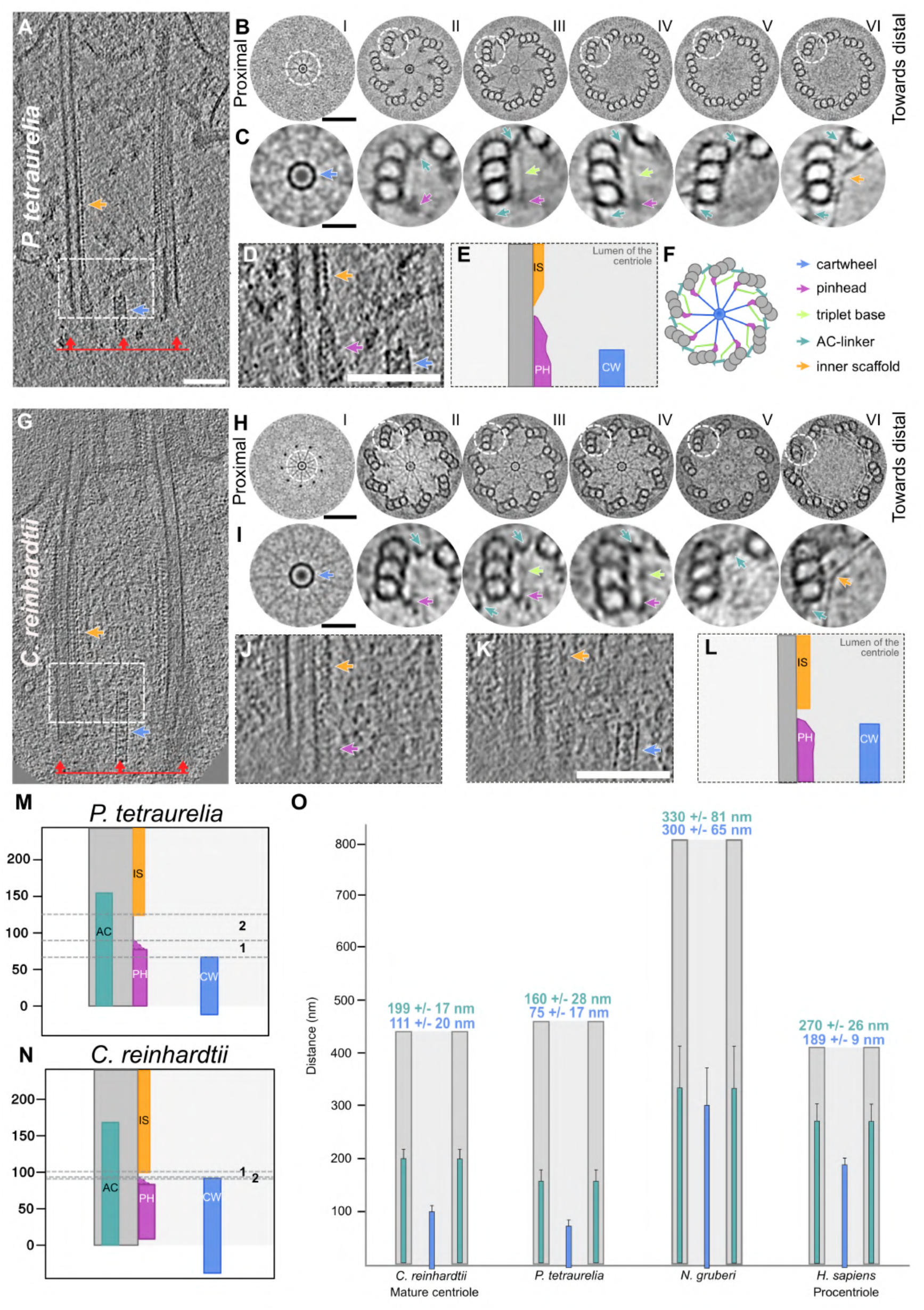
Structural features of the centriole’s proximal region in *P. tetraurelia* and *C. reinhardtii*. (A, G) Cryo-electron tomograms of *P. tetraurelia* (A) and *C. reinhardtii* (G) centrioles. Blue arrow denotes cartwheel, orange arrow denotes inner scaffold, red line with arrows denotes the proximal side of the centriole. Scale bar, 100 nm. (B, H) Nine-fold symmetrizations of serial cross sections taken along the proximal to distal axis in *P. tetraurelia* (B) and *C. reinhardtii* (H). Each section is a z-projection of 20.7 nm. White dashed circles delineate the structures highlighted in C and I. Scale bar, 60 nm. (C, I) Zoomed images of proximal centriole substructures from nine-fold symmetrizations of *P. tetraurelia* (C) and *C. reinhardtii* (I) along the proximal-distal axis. Each panel corresponds to the above image from panel B or H. Purple arrow, pinhead; light green arrow, triplet base; turquoise arrow, A-C linker; orange arrow, inner scaffold. (D, J, K) Side view showing the transition from pinhead to inner scaffold in *P. tetraurelia* (D) and *C. reinhardtii* (J, K). (E) Cartoon representation of panel D. (F) Representative model of a cross section of a centriole’s proximal region. Colored arrows indicate the different structural features identified. (L) Cartoon representation combining the z-projections in panels J and K. (M, N) Positioning of the different structures along the proximal length from representative *P. tetraurelia* (M) and *C. reinhardtii* (N) centrioles. Distance between the ends of the pinhead and cartwheel regions is denoted by zone 1 (for quantification, see Figure S7E). Distance between end of the pinhead region and start of the inner scaffold region is denoted by zone 2 (for quantification, see Figure S7F). (O) Cartwheel and A-C linker length in *C. reinhardtii, P. tetraurelia, N. gruberi*, and *H. sapiens*. Means and standard deviations of the mean are displayed above the range. A-C linker, turquoise; cartwheel, blue; microtubule triplets, grey.

On the basis of these observations, we measured the distance from the end of the pinhead region to the end of the cartwheel region and to the start of the inner scaffold in 5 *in situ C. reinhardtii* centrioles and 17 isolated *P. tetraurelia* centrioles. We found that the distances between these structural features is ∼5 nm on average in *C. reinhardtii*, which is close to the size of a tubulin monomer, indicating a direct transition from one structure to the other (Figure S7 E, F). In contrast, this gap distance is longer and more variable in *P. tetraurelia* centrioles, suggesting more stochasticity in the transitions between structures (Figure S7E, F). We also noted a strong correlation between the lengths of the A-C linker and the pinhead in *P. tetraurelia* centrioles (Figure S7B), suggesting that these two structures might have coordinated assembly. Conversely, there is no clear correlation between the lengths of the cartwheel and pinhead in *P. tetraurelia* centrioles (Figure S7C).

To better understand the relationship between the A-C linker and the cartwheel, we mapped their respective boundaries in the centrioles of *P. tetraurelia, C. reinhardtii, N. gruberi* and humans (Figure 4O). We found that the cartwheel length extends 111 +/− 20 nm, 75 +/− 17 nm, 300 +/− 65 nm and 189 +/− 9 nm in *C. reinhardtii, P. tetraurelia, N. gruberi* and humans, respectively (Figure 4O). Note that, as expected, mature human centrioles lacked cartwheels (Guichard et al., 2010), but we found 4 procentriole cartwheels to include in our analysis. In parallel, we analyzed the boundaries of the A-C linker and found that it spans 199 +/− 17 nm, 160 +/− 28 nm, 330 +/− 81 nm and 270 +/− 26 nm of the proximal region in *C. reinhardtii, P. tetraurelia, N. gruberi* and humans, respectively (Figure 4O). As previously reported (Le Guennec et al., 2020), this represents approximately 40% of the total centriole length. Comparing the measurements of these two structures reveals that the cartwheel spans 56% of the A-C linker length in *C. reinhardtii*, 47% in *P. tetraurelia*, 66% in *N. gruberi* and 70% in humans.

### The triplet base bridges the pinhead with the A-C linker

Our analysis of raw tomograms revealed that the triplet base emanates from the pinhead and binds the A-C linker, thereby indirectly connecting the cartwheel to the A-C linker (Figure 4). However, this analysis did not allow us to precisely detect where the triplet base connects to the A-C linker. Moreover, this connection has never been observed in previous subtomogram averaging analysis (Guichard et al., 2013; Li et al., 2019). Consequently, we undertook a subtomogram averaging approach focused on revealing the triplet base connection and the A-C linker structure, using 11 tomograms of uncompressed *P. tetraurelia* centrioles. We succeeded in resolving the triplet base in our average; however, it had very low map density, suggesting that this structure is flexible or not stoichiometrically occupied (Figure 5A) and explaining why it has not been observed before in cryo-ET. It is also important to note that although both the triplet base and the pinhead are clearly visible, we could not reliably retrieve their longitudinal periodicities.

**Figure 5.**
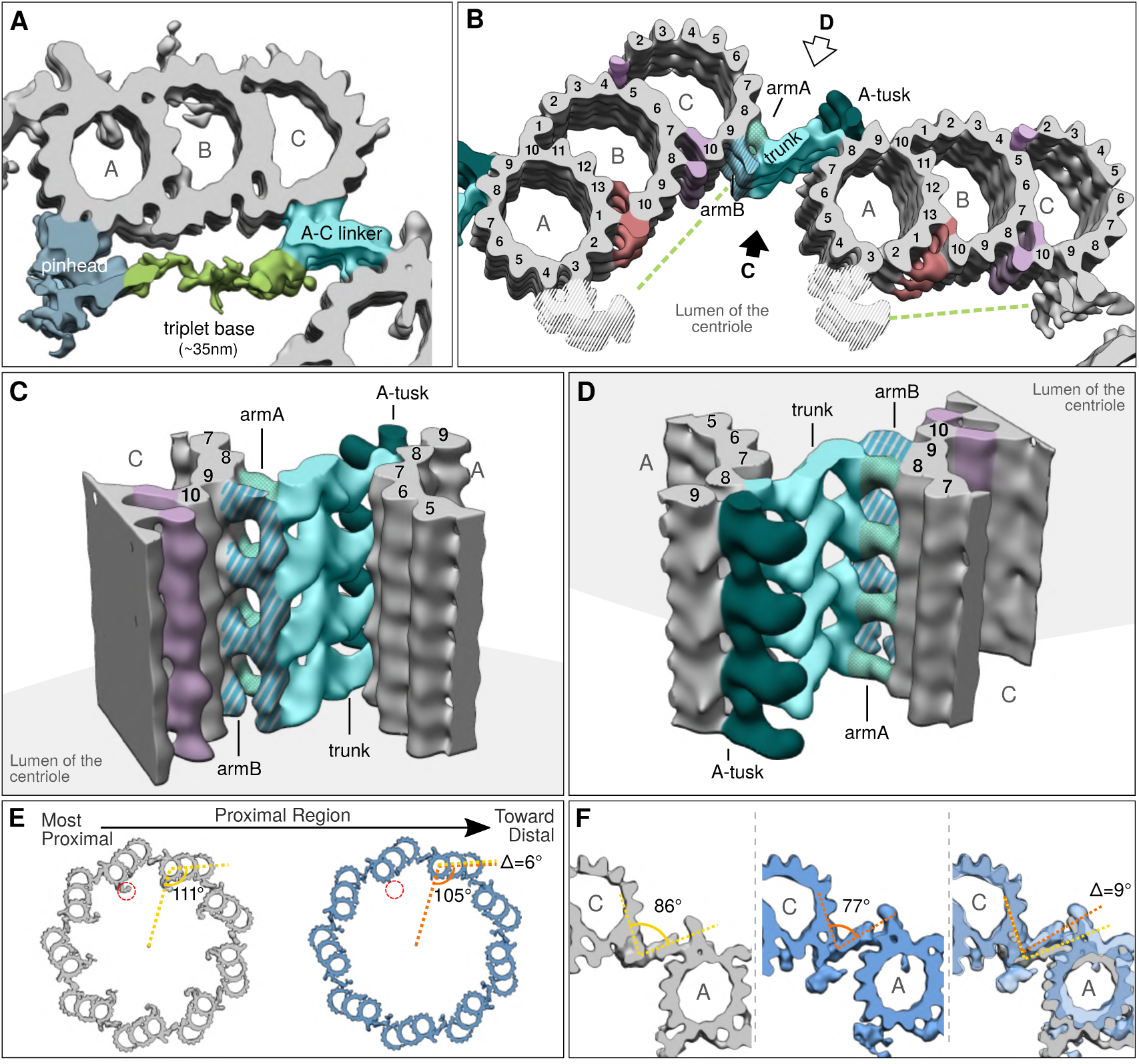
Subtomogram averaging of the proximal triplet from *P. tetraurelia*. (A) Microtubule triplet reconstruction from the beginning of the proximal region, displayed with a low contour threshold value to show the triplet base density (green) connected to the pinhead (blue) and the A-C linker (turquoise). (B) Two adjacent triplets from the beginning of proximal region, displayed with a higher contour threshold than in A. The A-C linker is segmented into different substructures (patterned turquoise colors) according to nomenclature (Li et al., 2019). The green dashed line indicates the putative position of the triplet base. Non-tubulin densities are colored in red and purple. The pinhead has been hidden in this view, as its reconstruction is not correct due to the 8.5 nm initial subvolume picking that imposes this periodicity on the structure. (C) Three-dimensional side view of the A-C linker from the lumen of the centriole. (D) Three-dimensional side view of the A-C linker from outside the centriole (rotated 180° from C). (E) Top views of independent averages from the more proximal (grey) and more distal (blue) parts of the *P. tetraurelia* proximal region. (F) Focus on the A-C linker from the beginning of the proximal region (left, grey), the end of the proximal region (middle, blue), and the superimposition of both structures (right).

Next, we focused on the A-C linker and found that it can be subdivided into two major regions previously observed in *Trichonympha*: the A-link that contacts the A-tubule and the C-link that contacts the C-tubule. The *P. tetraurelia* A-C linker has a longitudinal periodicity of 8.4 +/− 0.2 nm, consistent with previous measurements from *Trichonympha* and *C. reinhardtii* (Guichard et al., 2013; Li et al., 2019) (Figure S8A, B and Figure S8F-H). With the obtained resolution of 31.5Å (Figure S9), we were able to identify that the C-link is composed of two main densities: ArmA, which contacts the C-tubule protofilaments C8 and C9, and ArmB, which decorates only C-tubule protofilament C9 (Figures 5 and S8). On the A-link side, we identified a single connection between the A-link’s trunk and A-tubule protofilament 8, an interaction originally described in *C. reinhardtii* (Li et al., 2019) (Figure S8F, G). In addition to the A-C linker, we identified a large density between protofilaments 8 and 9 of the A-tubule that we termed the A-tusk (Figures 5 and S8C-E). Interestingly, we observed that the triplet base connects to the A-C linker directly on the ArmB density, reinforcing our conclusion that the entire proximal region forms an interconnected structural network from the central hub of the cartwheel, through the radial spokes, the pinhead and the triplet base to the A-C linker.

To check whether the connection between the pinhead and the A-C linker is maintained throughout the proximal region, we split the dataset in two halves corresponding to the more proximal and more distal parts of this region (Figure 5E, F). The nine-fold symmetrized model of each map was reconstructed. Interestingly, as previously observed (Figure 4 B, C), we noticed that the pinhead density is almost completely absent in the average from the more distal part of the proximal region, whereas the A-C linker is still present and has an extra density on ArmB seemingly replacing the triplet base position (Figure 5E, F, red circles). This observation indicates that although the pinhead and A-C linker are connected through the triplet base, the presence of the A-C linker is independent of the pinhead and triplet base. We also noticed a difference in the microtubule triplet and A-C linker angles between the two maps (Figure 5F), with an angle decrease of 6° for the triplet and 9° for the A-C linker. As this difference was previously observed in *C. reinhardtii* (Li et al., 2019), the slight twist we measured in the proximal region appears to be evolutionarily conserved. This proximal twist suggests that the A-C linker is able to adapt to the difference in angles between the microtubule triplets and thus remain connected to them.

## Discussion

In this study, we used cryo-ET to analyze the proximal region of centrioles from four evolutionarily distant species. We describe the structural features of this region including the cartwheel, the pinhead, the triplet base and the A-C linker (Figure 6). Interestingly, we found that the cartwheel structure protrudes proximally beyond the microtubule triplets in all species that we investigated, especially in the assembling *C. reinhardtii* procentrioles. This observation supports the notion that the cartwheel assembles independently of the microtubule triplets, which are connected by the A-C linker, and that the two structures, cartwheel and A-C linker, likely play a role in defining the nine-fold symmetry of the organelle (Hilbert et al., 2016; Nakazawa et al., 2007) as well as its cohesion at the proximal region (Le Guennec et al., 2020; Yoshiba et al., 2019). The cartwheel’s proximal extension is also consistent with the proximal-directed growth of the cartwheel protein SAS-6 observed in *Drosophila* (Aydogan et al., 2018). It is currently not known whether the cartwheel structure can grow from its proximal end and whether such a mechanism is evolutionary conserved.

**Figure 6.**
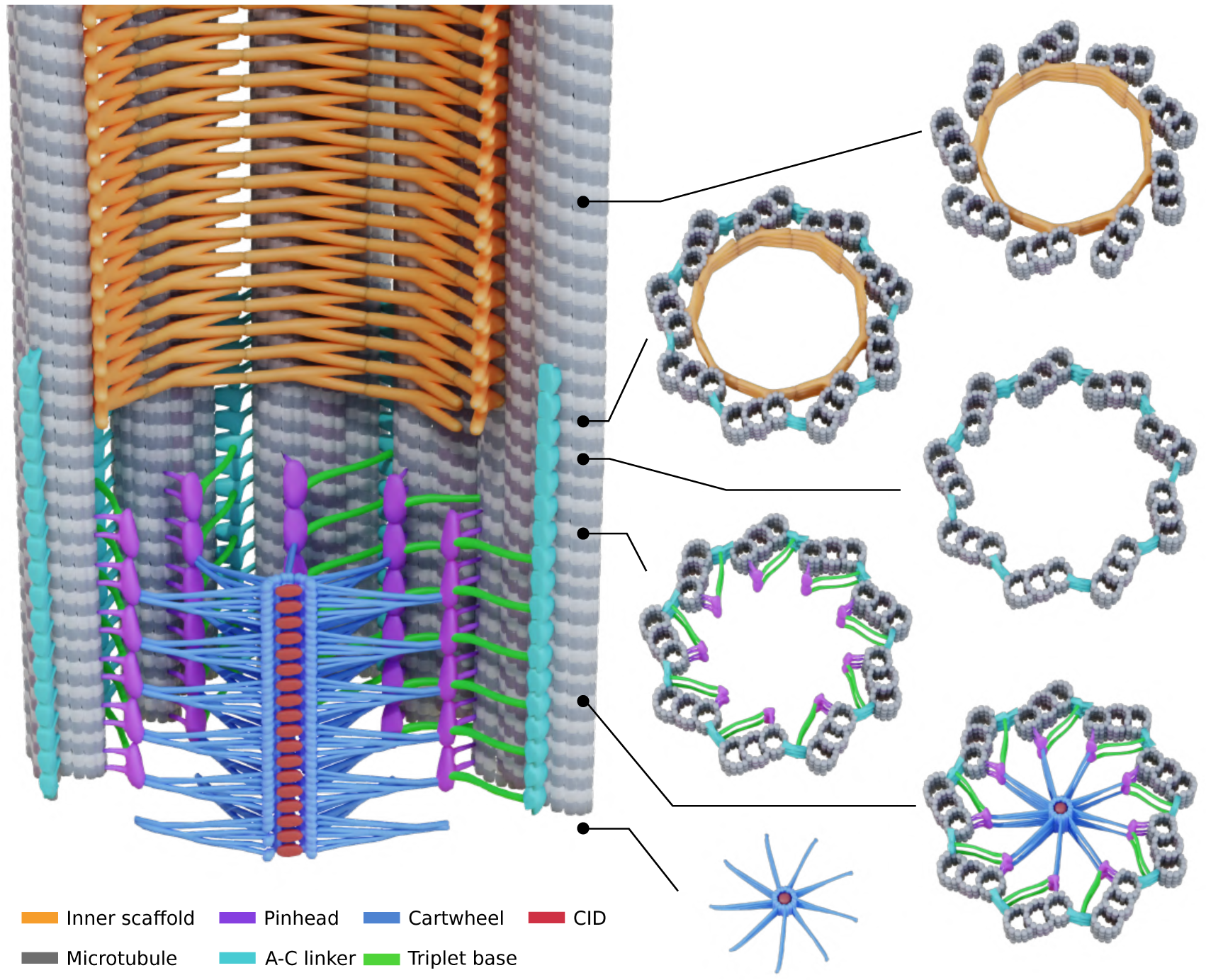
Model of the architecture of the proximal region of the centriole. The colors corresponding to each structure are indicated in the legend. Note that the cartwheel structure protrudes proximally from the microtubule wall; here, one unit has been depicted that corresponds to an external cartwheel of about 25nm. The cartwheel’s structural unit consists of 3 ring-pairs, from which emanate 6 radial spokes that merge into one density before contacting the pinhead structure. The pinhead and the A-C linker are connected through the triplet base. The A-C linker extends more distal than the cartwheel and co-exists with the inner scaffold structure.

Our cryo-ET analysis revealed that the cartwheel’s central hub in *C. reinhardtii* and *P. tetraurelia* is organized in ring-pairs (Figure 6). Furthermore, in all four studied species, we observed densities inside the lumen of the central hub with a similar periodicity formed by the CIDs in *Trichonympha*. We therefore conclude that CID structures are present in every species studied to date and are a conserved element of the cartwheel. Moreover, one CID is positioned between the two rings of the ring-pair, suggesting that it might be involved in ring-pair assembly by helping build a cohesive unit. Whether the ring-pair is composed of only SAS-6, or whether another protein participates in forming this structure, is an open question that the resolution of our current dataset cannot answer. Therefore, an important future challenge will be to determine the molecular composition of the ring-pair and the CID.

At the outer margin of the central hub’s ring-pairs, we observed that the cartwheel spokes are clearly organized differently than in *Trichonympha*, which turns out to be the biggest structural difference between the cartwheels of the different species. In *Trichonympha*, we could observe only two spokes merging, forming a longitudinal periodicity of 17 nm (Guichard et al., 2013). Here, we have demonstrated that the resolution obtained in the *Trichonympha* study is not sufficient to see certain details. Nevertheless, even by artificially lowering the resolution of our *P. tetraurelia* cartwheel map, the spoke organization remains distinct, with a lateral periodicity of ∼25 nm. In both *C. reinhardtii* and *P. tetraurelia* cartwheels, this 25 nm periodicity results from the merge of spokes emanating from 3 adjacent ring-pairs (Figure 6). However, we could also distinguish that the spoke organization differed between these species. In *P. tetraurelia*, one spoke is made of 3 substructures that each emanate from a pair of rings, whereas in *C. reinhardtii*, the final spoke tip is made from only two substructures (Figure 3J). As the coiled-coil domain of SAS-6 is part of the spokes (Gönczy, 2012), the difference in radial spoke organization could potentially be explained by the low homology between SAS-6 coiled coils (Leidel et al., 2005). It is possible to imagine that coiled coils of neighboring SAS-6 proteins merge to form a coiled coil bundle or a tetramer/hexamer. Another possibility is that a different protein interacts with the SAS-6 coiled coil and is responsible for this bundling. To date, SAS-5 is one of the most likely candidates for this role. Indeed, it has been shown in several species that SAS-5 interacts with the SAS-6 coiled coil where the bundle is formed (Cottee et al., 2013; Qiao et al., 2012; Shimanovskaya et al., 2013). In addition, it has been shown that the Ana2 (SAS-5 in *Drosophila*) coiled coil forms a tetramer (Cottee et al., 2013) and that *C. elegans* SAS-5 forms higher-order protein assemblies up to hexamers in solution (Rogala et al., 2015). It is therefore possible that different stoichiometries of SAS-6:SAS-5 can modify the architecture of the spoke bundling.

Our study also highlights the triplet base structure (Figure 6), originally described in conventional electron microscopy of resin-embedded mammalian centrioles (Vorobjev and Chentsov, 1980). We found that the triplet base connects the pinhead to the A-C linker, thus forming a continuous structure that bridges the cartwheel with the A-C linker. The triplet base might enhance the cohesion and stability of the entire proximal region. Although its molecular nature is not known, its apparent flexibility, length and low map density, similar to the cartwheel spokes, would suggest that the triplet base is made by a long coiled coil protein. It is therefore tempting to speculate that this structure might consist of the coiled coil protein Bld10p/Cep135. Indeed, based on its immuno-localization as well as its known interaction with the C-terminus of SAS-6 and microtubules, current models place this protein as part of the pinhead (Hiraki et al., 2007b; Hirono, 2014; Kraatz et al., 2016). The coiled coil length prediction for Cep135 is ∼900 of its 1140 total amino acids, which would yield a coiled coil that is 133 nm long (900 residues x 0.1485 nm [axial rise per residue]= 133 nm, formula from (Kitagawa et al., 2011)). Considering that the pinhead is ∼20 nm long (Guichard et al., 2013), it is likely that a large portion of Bld10/Cep135 extends from it. Therefore, we propose that a part of the predicted 133 nm coiled coil constitutes the 35 nm long triplet base connecting to the A-C linker (Figure 5A). This hypothesis is consistent with the phenotypes of *C. reinhardtii* and *Tetrahymena* Bld10p mutants, which not only lose the connection of the cartwheel to the microtubule wall but also lose the microtubule triplets themselves, suggesting that the cohesion between triplets is partially lost (Bayless et al., 2012; Matsuura et al., 2004). Future studies on the precise location of the different regions of Cep135 would be needed to answer these questions.

An important structural feature revealed in our study is the intrinsic polarity of the cartwheel structure along its proximal-distal axis. Previous work had observed such polarity in the pinhead and A-C linker structure (Guichard et al., 2013; Li et al., 2019). Our work now reveals that polarity also exists within the cartwheel itself, which might play a critical role in centriole biogenesis. Such polarity is likely important to define the directionality of structural features that assemble after cartwheel formation. For instance, microtubule triplets, which are also polarized structures, only grow in the distal direction. Although it is possible that the triplets lengthen slightly on the proximal side, it is clear that the plus ends of the microtubules always face the distal end of the centriole. It is therefore possible that the polarity of the cartwheel defines the growth directionality of the procentriole from the very beginning of assembly. It is interesting to note that the only known example of microtubule triplet polarity inversion was observed in a *Tetrahymena* Bld10p mutant (Bayless et al., 2012). As Bld10p constitutes part of the cartwheel spoke-tip/pinhead, this reinforces the idea that the cartwheel defines the direction of centriole growth.

Combining our present study with previous work on the structure of the centriole proximal region from different species offers a glimpse at evolutionary conservation and divergence at the level of molecular architecture. The data presented here suggest that the cartwheel-containing region has a conserved overall organization with defined structural characteristics (Figure 6). However, our work also demonstrates that the specific layout of the centriole and the finer structural elements may differ considerably between species. These observations correlate well with the fact that many centriolar proteins are conserved between species, yet they can vary significantly in their size or amino acid composition, as exemplified by the low sequence homology of the cartwheel protein SAS-5/Ana2/STIL (Stevens et al., 2010). Our work therefore shows that there may be different routes to build a centriole.

## Author Contributions

V.H, P.G. and B.D.E conceived, supervised, designed the project and wrote the final manuscript with input from all authors. N. K and M.L.G. performed all image processing and analyzed the data. A-M.T. purified the *P. tetraurelia* centrioles. N. K. isolated the human centrioles and acquired tomograms of these two species with the help of L.K, K.N.G., H.v.d.H and B.D.E. Sample preparation and tomography of in situ *C. reinhardtii* centrioles as well as isolated *N. gruberi* centrioles was performed by P.E., M.S., H.v.d.H and B.D.E. G.A generated the 3D model of the centriole.

## Acknowledgments

We thank the BioImaging Center at Unige. We thank Jürgen Plitzko and Wolfgang Baumeister for providing support and instrumentation. This work was supported by the Swiss National Science Foundation (SNSF) PP00P3_187198 and by the European Research Council ERC ACCENT StG 715289 attributed to P.G., as well as the Helmholtz Zentrum München and the Max Plank Society.

## Competing Interests

The authors declare that they have no competing interests.

## Data and materials availability

Subtomogram averages have been deposited at the Electron Microscopy Data Bank (EMD-10726, EMD-10727, EMD-10728, EMD-10729). All data needed to evaluate the conclusions in the paper are present in the paper and/or the supplementary materials. Additional data is available from authors upon request. Correspondence and requests for materials should be addressed to P.G.

---

**Figure S1.**
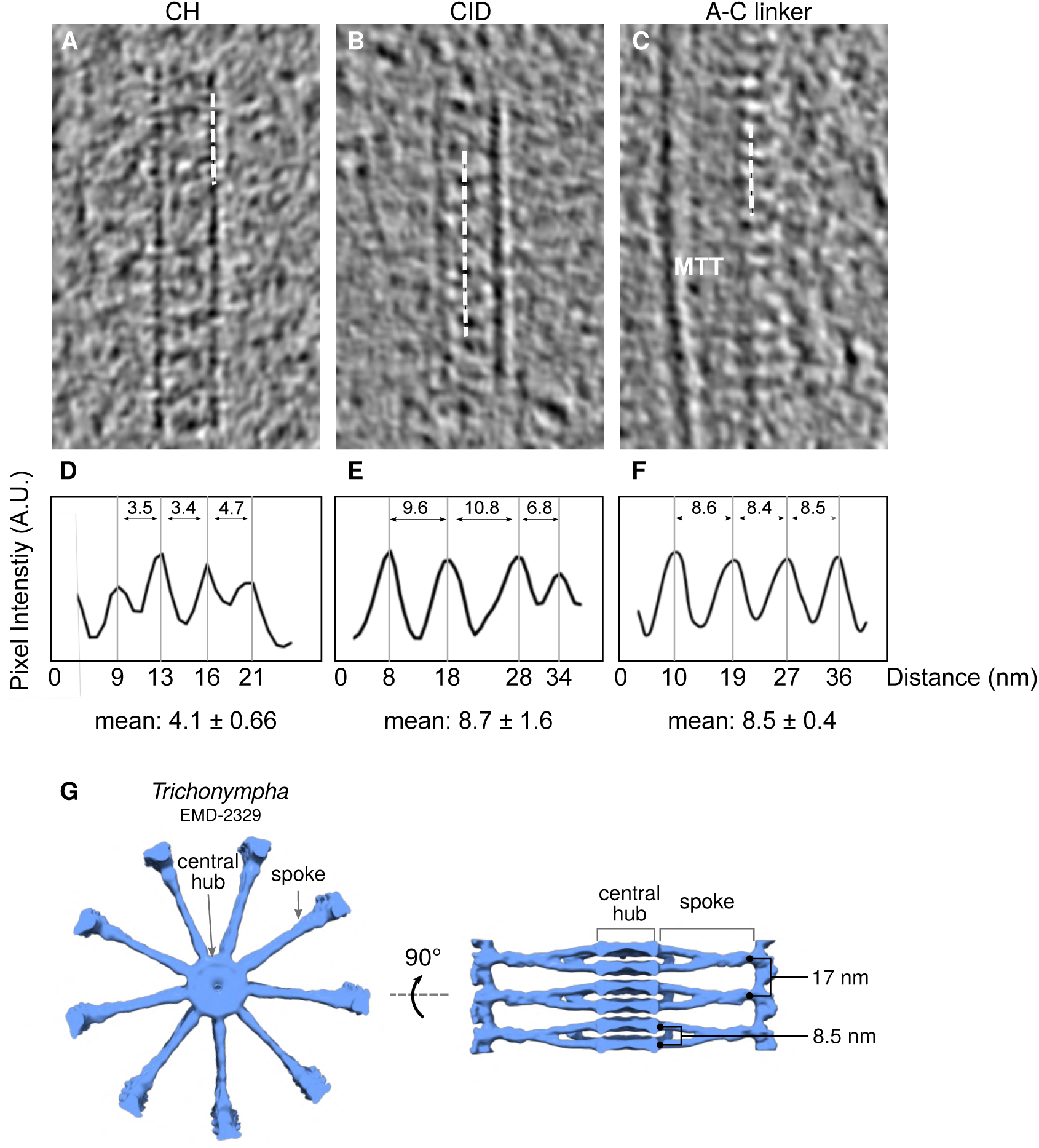
Periodicity along the central hub, cartwheel inner densities, and A-C linker in *C. reinhardtii in situ* centrioles. (A, B, C) Cryo-ET sections depicting representative central hub (CH) (A), several cartwheel inner densities (CIDs) (B), and A-C linker (C). Dashed white line denotes region from which plot profiles were generated. (D, E, F) Plot profiles with their associated mean periodicity displayed below. Scale bar, 20 nm. (G) Top and side views of *Trichonympha* cartwheel and associated periodicities from (Guichard et al., 2013).

**Figure S2.**
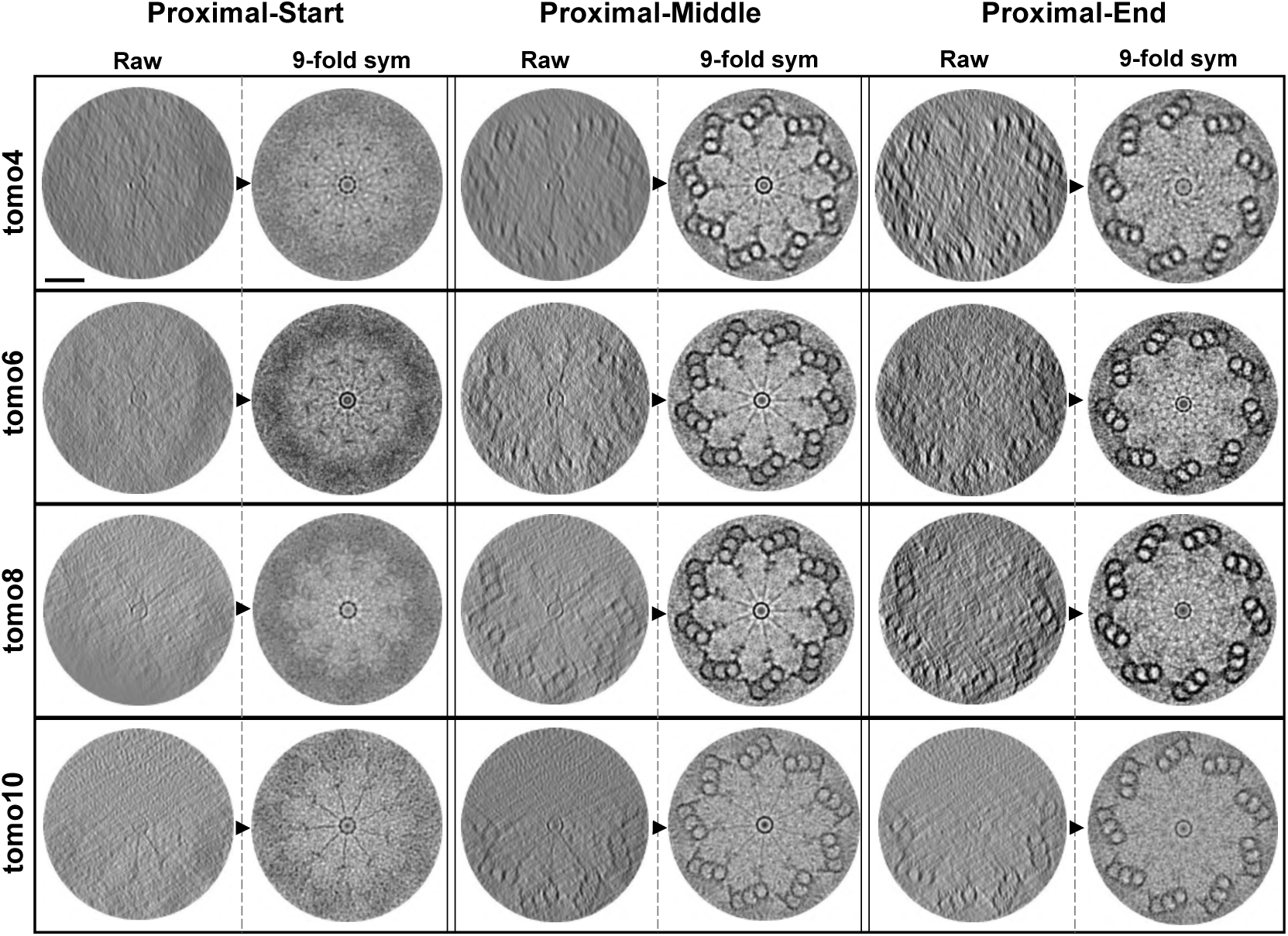
The cartwheel-containing region in mature *C. reinhardtii* centrioles. From four tomograms, three regions were extracted every 35 nm along the proximal region corresponding to the Proximal-Start, Proximal-Middle, and Proximal-End regions. Each image corresponds to a projection of about 27 nm. To improve visualization, we applied a nine-fold symmetrization of the image (displayed to the right of the raw image, separated by a dashed grey line and a black arrowhead). Scale bar, 100 nm.

**Figure S3.**
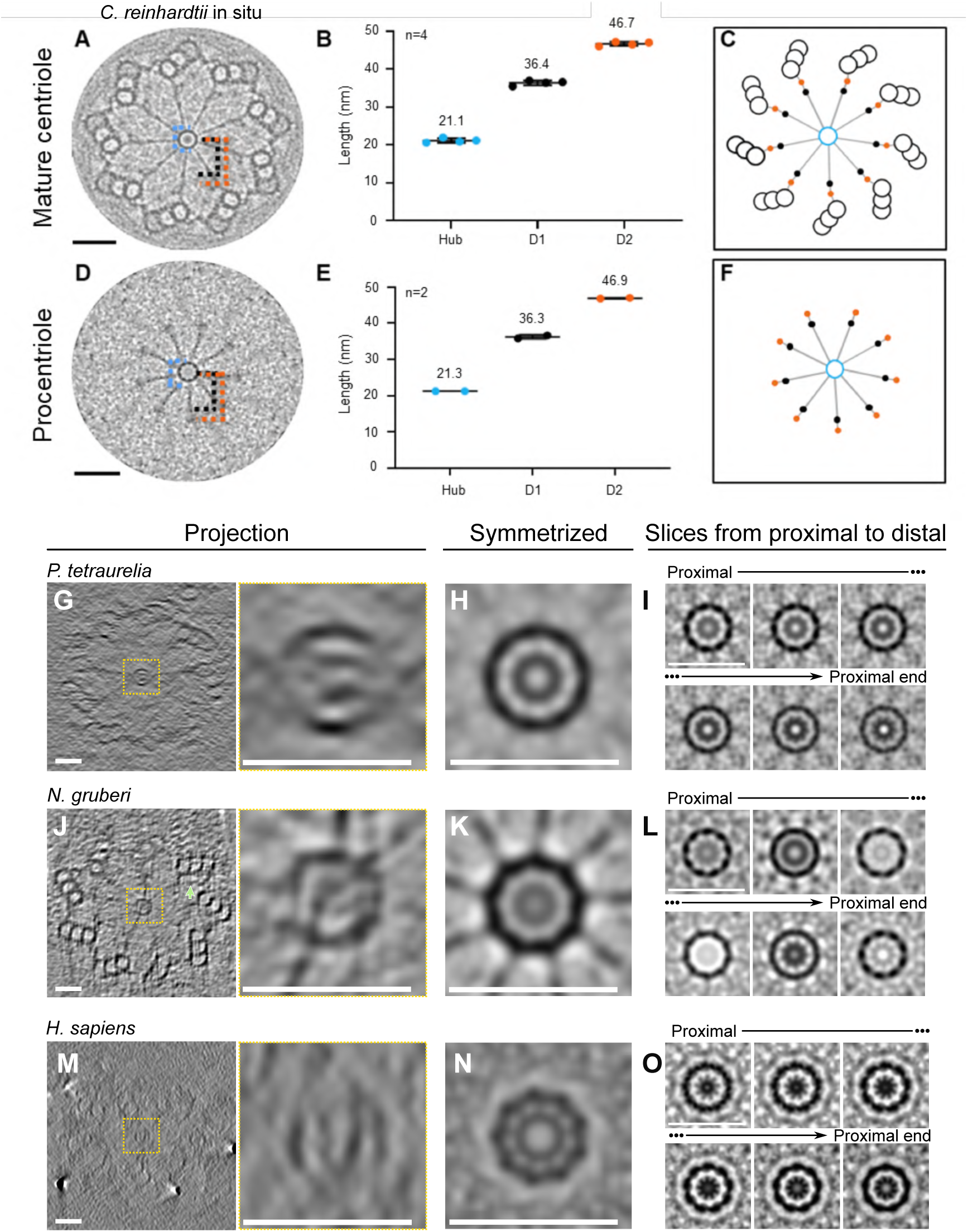
*In situ* spoke architecture of the *C. reinhardtii* cartwheel, and conservation of the cartwheel inner densities. (A, D) Nine-fold symmetrized cross sections of cartwheel-containing regions from a mature centriole (A) and a procentriole (D), both from *in situ* tomograms of *C. reinhardtii*. The dashed blue line indicates central hub diameter. Dashed black and orange lines indicate distances from the external edge the central hub to D1 and D2 densities of the radial spoke, respectively. (B, E) Measurements of cartwheel features: mean diameter of the central hub (blue), distance from the central hub to D1 (black) and D2 (orange) densities in mature centrioles (B, n = 4) and procentrioles (E, n = 2). Mean values are displayed above data range. (C, F) Models of cartwheel organization and distance from the central hub to D1 and D2 in mature centrioles (C) and procentrioles (E). Central hub, blue; spoke, grey; D1, black circle; D2, orange circle. (G, J, M) Cryo-electron tomogram cross sections depicting top views of the proximal regions of *P. tetraurelia* (G), *N. gruberi* (J), and *H. sapiens* (M). Yellow dashed box indicates central hub with the corresponding zoom on the right. Scale bar, 40 nm. (H, K, N) Nine-fold symmetrized images corresponding to the panels in A, J and M. (I, L, O) Symmetrized 4 nm serial projections through one central hub of *P. tetraurelia* (I), *N. gruberi* (L), and *H. sapiens* (O). Scale bar, 40 nm.

**Figure S4.**
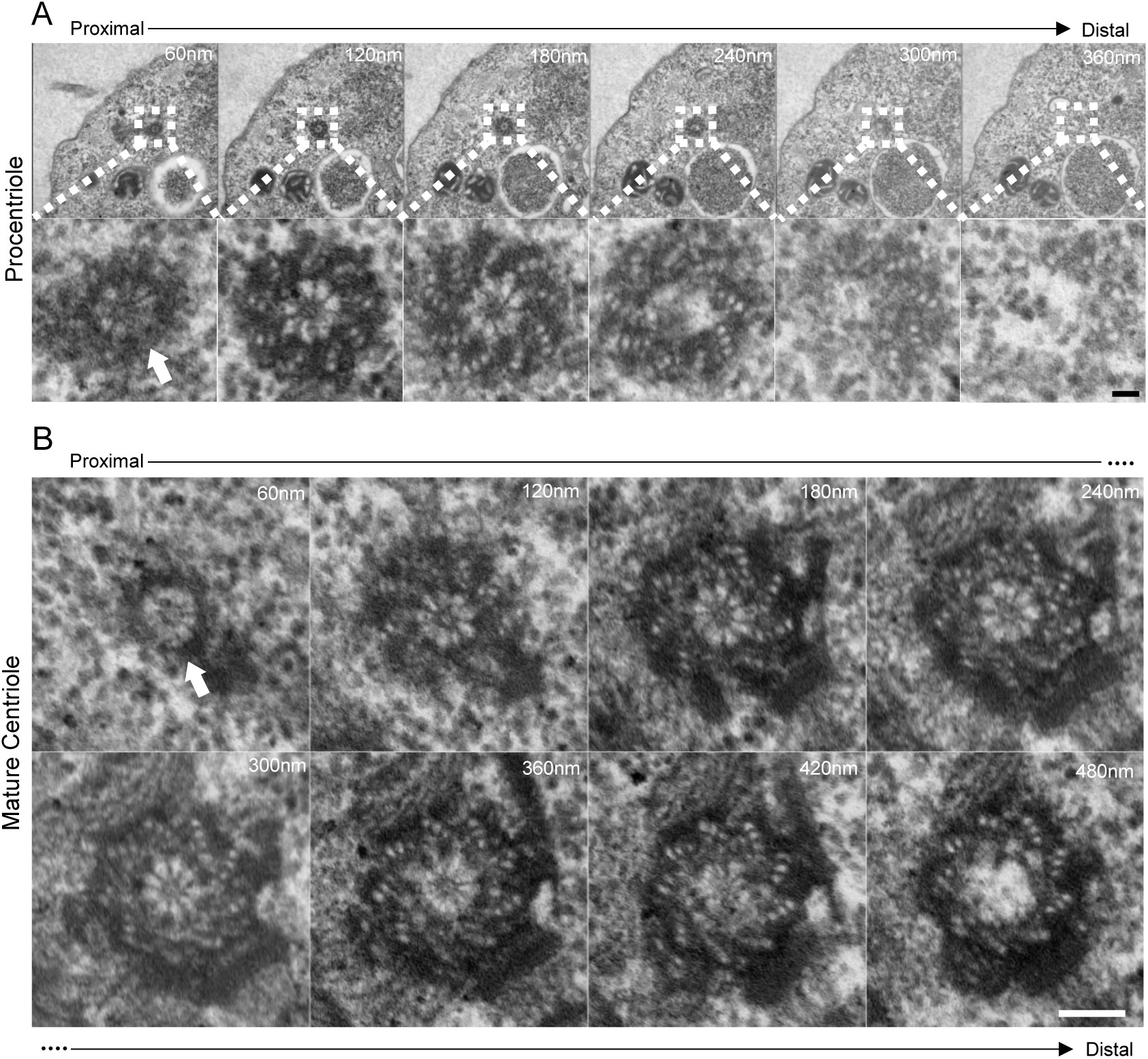
Resin-embedded *N. gruberi* cells display the proximal cartwheel protrusion in both mature centrioles and procentrioles. (A) Serial sections through a procentriole in a resin-embedded *N. gruberi* cell, moving from proximal (left) to distal (right). White-dashed box denotes the zoomed region in the bottom panel. White arrow denotes the cartwheel extending beyond the proximal microtubule triplet region. Scale bar, 50nm. (B) Serial sections through a mature centriole in a resin-embedded *N. gruberi* cell, moving from proximal (left) to more distal (right). White arrow denotes the cartwheel extending beyond the proximal microtubule triplet region. Scale bar, 100nm

**Figure S5.**
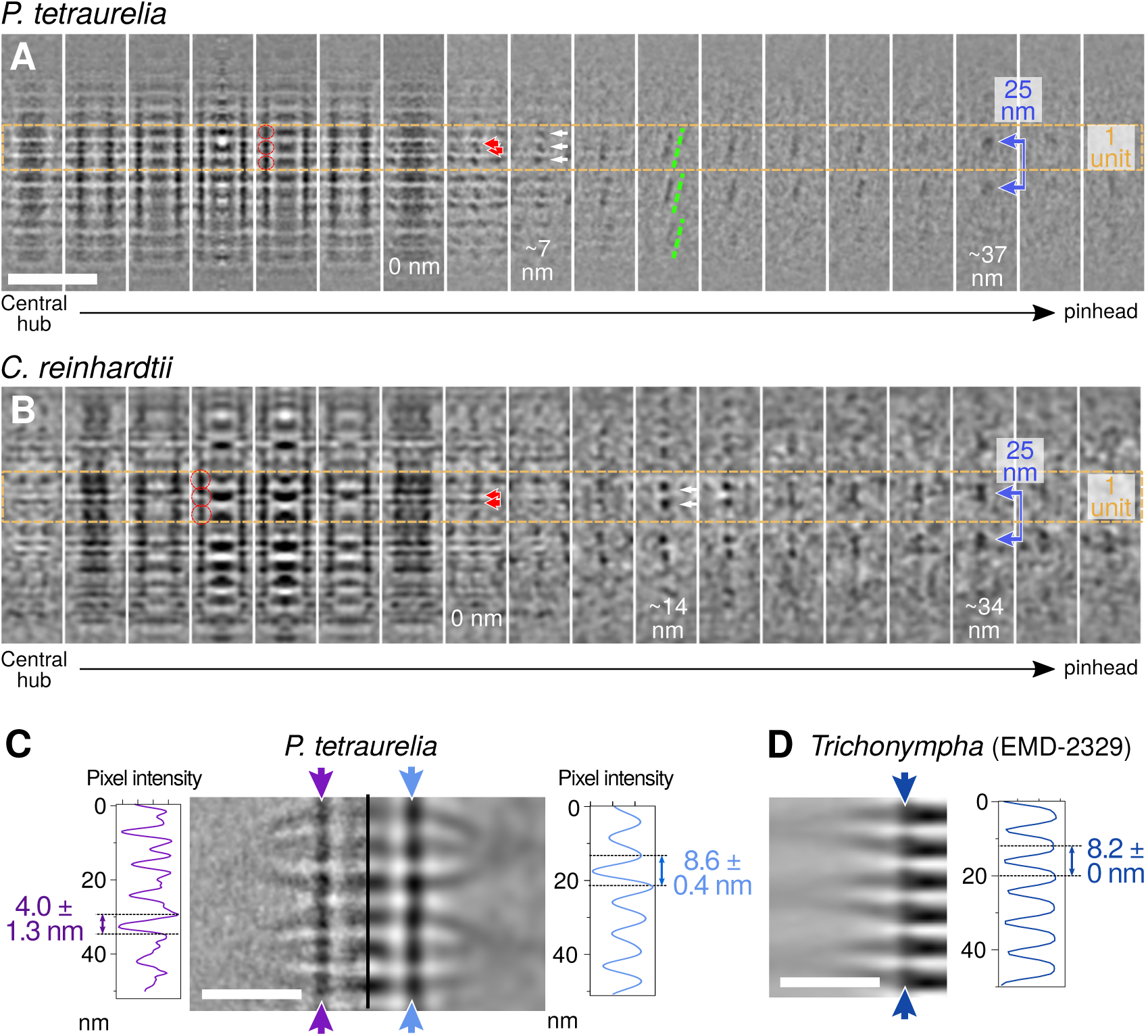
Cartwheel spoke organization in *P. tetraurelia* and *C. reinhardtii* from the central hub through the pinhead. (A, B) Serial z-projections of approximately 4 nm thickness from subtomogram averages of *P. tetraurelia* (A) and *C. reinhardtii* (B) cartwheels. The left-most z-projections display the central hub, the right-most projections show the pinhead. Orange dashed lines delineate one repeat unit of the cartwheel. Red arrows mark individual spokes, white arrows mark merged spokes, and blue arrows with a line mark the final merged spoke (D1 density) longitudinally spaced every 25 nm. Scale bars, 50 nm. (C) Bandpass filter applied to a *P. tetraurelia* subtomogram average projection with a cutoff at 38 Å. Purple and blue arrows denote the central hub and position of the associated plot profiles. The unfiltered projection displays a mean periodicity of 4.0 +/− 1.3 nm (SEM), while the projection filtered to 38 Å displays a mean periodicity of 8.6 +/− 0.4 nm (SEM). Scale bar, 20nm. (D) Plot profile along the previously published *Trichonympha* central hub (EMD-2329) displaying a longitudinal periodicity of 8.2 nm. Scale bar, 20nm.

**Figure S6.**
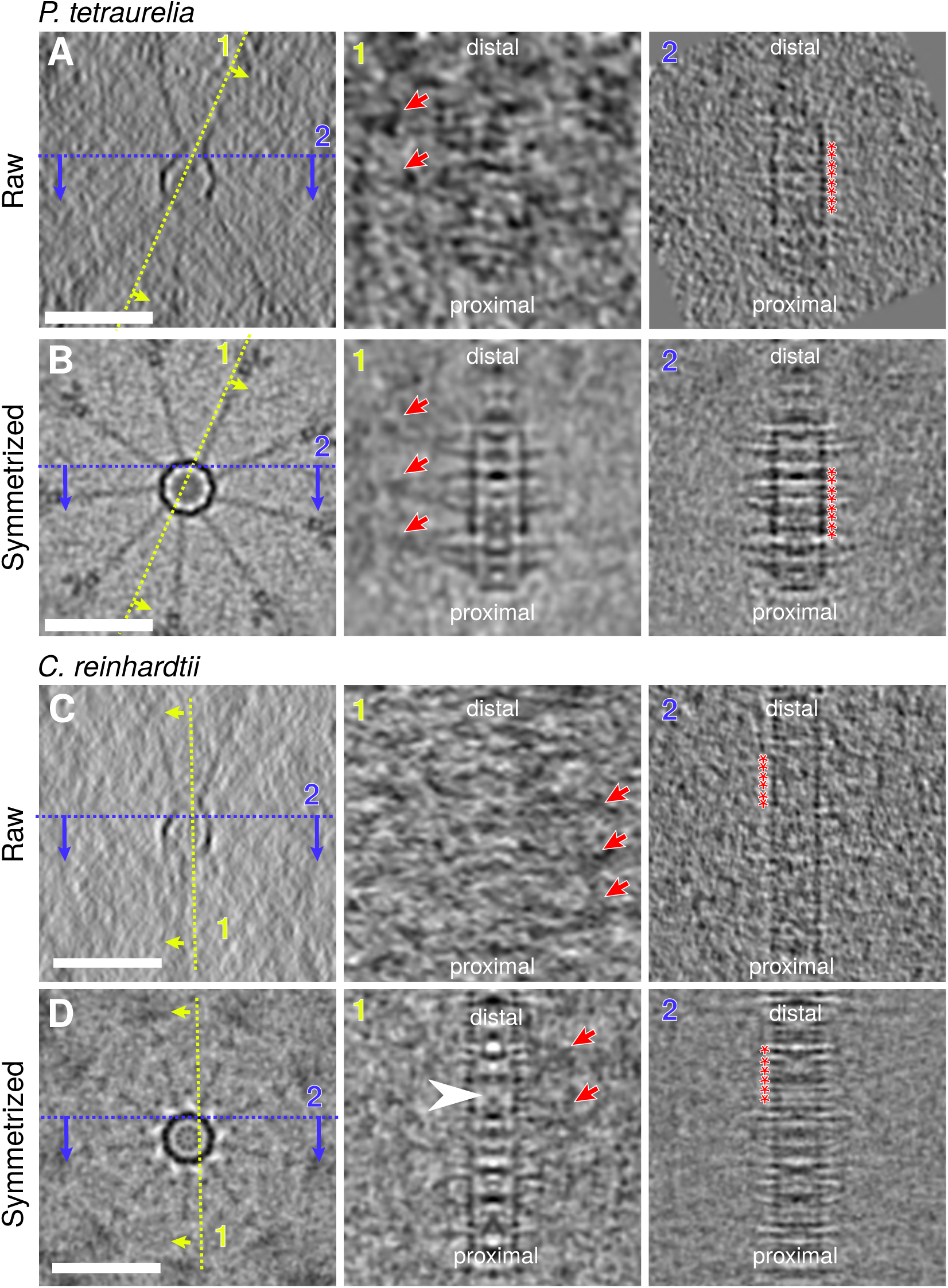
Raw and symmetrized cartwheels from *P. tetraurelia* and *C. reinhardtii*. (A, C) Cryo-electron tomogram sections displaying the cartwheels of *P. tetraurelia* (A) and *C. reinhardtii* (C) from top view (left panels) and side views (middle and right panels). Scale bars, 50 nm. (B, D) Corresponding nine-fold symmetrized image displaying the cartwheels *of P. tetraurelia* (B) and *C. reinhardtii* (D) from top view (left panels) and side views (middle and right panels). Scale bars, 50 nm. Dashed yellow lines and arrows indicate the position and direction of the reslice to visualize the radial spokes (1), Dashed blue line and arrows indicate the position and direction of the reslice to visualize the central hub (2). Red arrows indicate the position of merged spokes (1). Red asterisks denote positions of central hub ring subunits (2).

**Figure S7.**
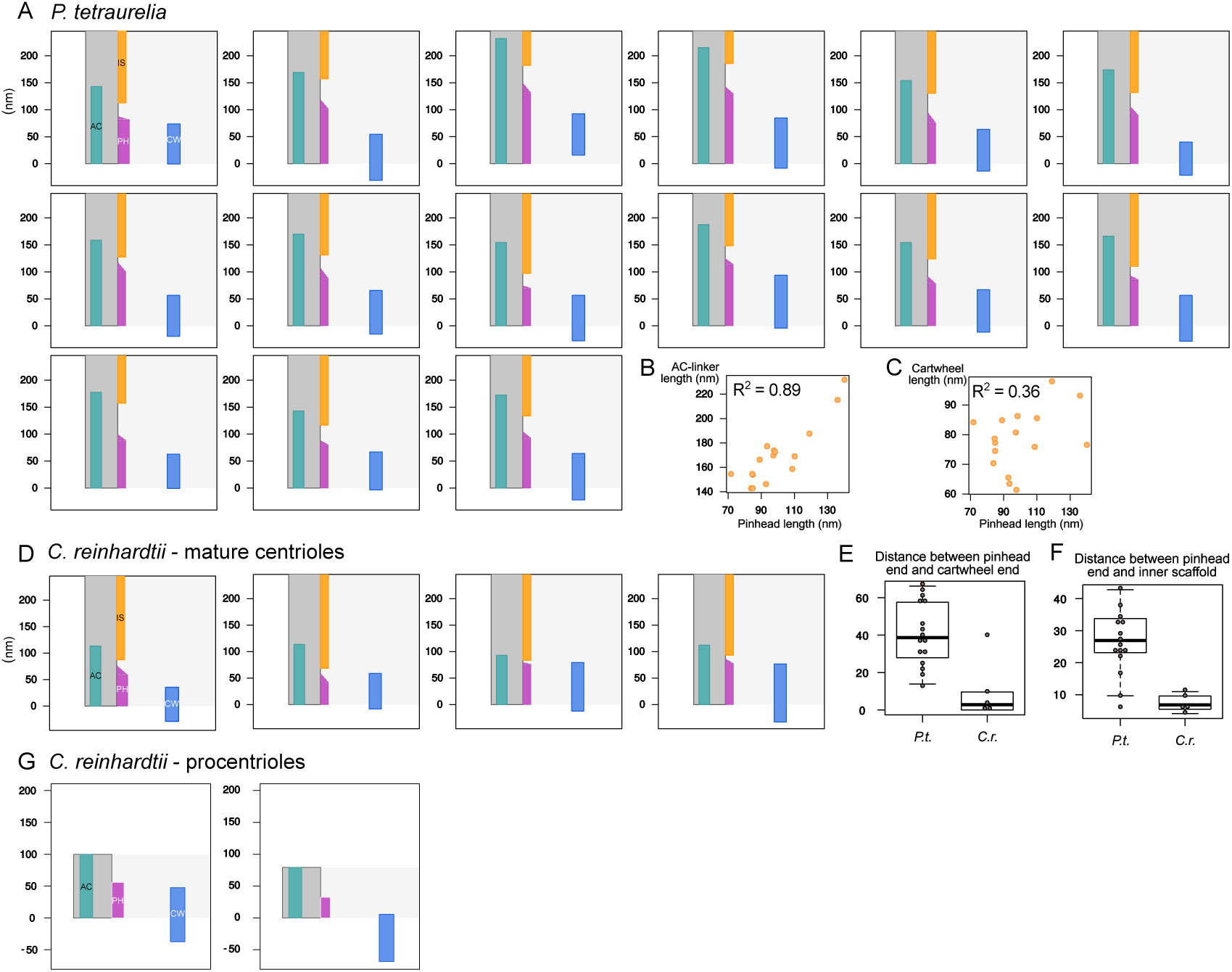
Boundaries of the proximal region’s structural features in *P. tetraurelia* and *C. reinhardtii* centrioles. (A) Position of the different structures along the proximal to distal axis of 15 different *P. tetraurelia* centrioles. Dark blue, cartwheel (CW); magenta, pinhead (PH); turquoise, A-C linker (AC); orange, inner scaffold (IS); dark grey, microtubules wall. (B) Correlation plot depicting A-C linker length versus pinhead length from *P. tetraurelia* centrioles. N = 16, Pearson correlation coefficient 0.89. (C) Correlation plot depicting cartwheel length versus pinhead length from *P. tetraurelia* centrioles. N = 16, Pearson correlation coefficient 0.36. (D) Position of the different structures along the proximal to distal axis of 4 different *C. reinhardtii* mature centrioles. Same color code as panel A. (E) Distance between the end of the pinhead region and the end of the cartwheel region. (n = 16, *P. tetraurelia*; n = 5, *C. reinhardtii*). (F) Distance between the end of the pinhead region and the beginning of the inner scaffold region (n = 15, *P. tetraurelia*; n = 5, *C. reinhardtii*). (G) Position of the different structures along the proximal to distal axis of 2 different *C. reinhardtii* procentrioles.

**Figure S8.**
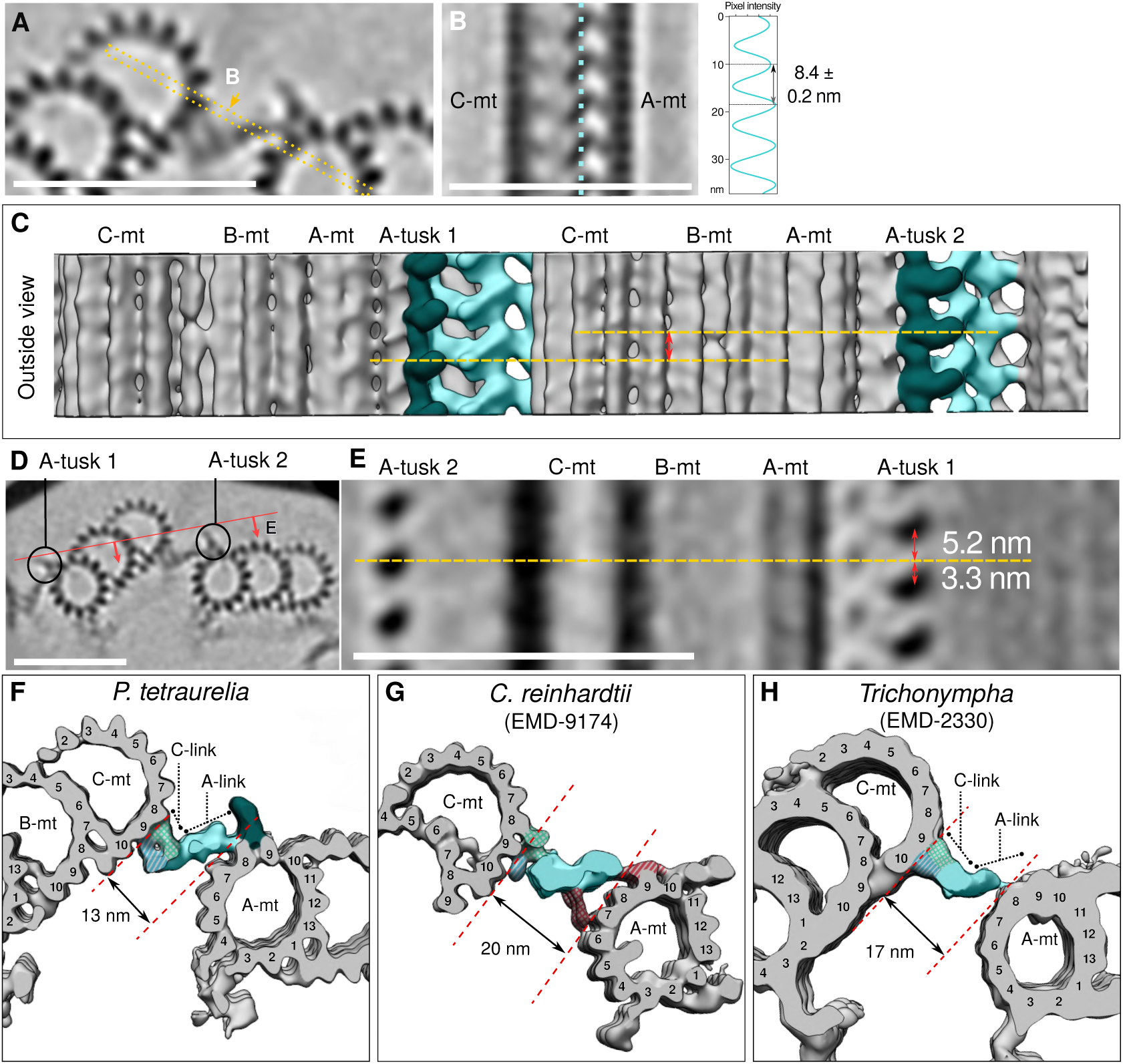
Architectural features of the *P. tetraurelia* proximal region, and evolutionary comparison of the A-C linker. (A) Z-projection of the reconstructed junction between adjacent proximal microtubule triplets. Scale bar, 50 nm. (B) Cross-section highlighting the lateral periodicity of the trunk and its associated plot profile (right) measured along the light blue dotted line. Scale bar, 50 nm. (C) Three-dimensional view of two adjacent proximal microtubule triplets seen from the outside of the centriole. Yellow dashed lines indicate the position of the A-tusk of adjacent triplets. The double headed red arrow indicates the shift along the z-axis between the position of two consecutive A-tusks. (D) Z-projection image of two adjacent proximal microtubule triplets. The red line indicates the position of the cross-section shown in (E). Scale bar, 50 nm. (E) Cross-section of two proximal microtubules triplets showing the shift along the z-axis of the A-tusk on one triplet (A-tusk 2) compared to the A-tusk on the adjacent triplet (A-tusk 1). Scale bar, 50 nm. (F, G, H) Three-dimensional views of *P. tetraurelia* (F), *C. reinhardtii* (G, EMD-9174, filtered to 45 Å) and *Trichonympha* (H, EMD-2330). The dotted red lines define the distance between consecutive microtubule triplets. Note that this distance varies between species. Microtubule triplets are in grey and the A-C linker is in light blue/green. Dashed blue: arms A and B, blue: trunk, red: legs. Dark green: A-tusk.

**Figure S9.**
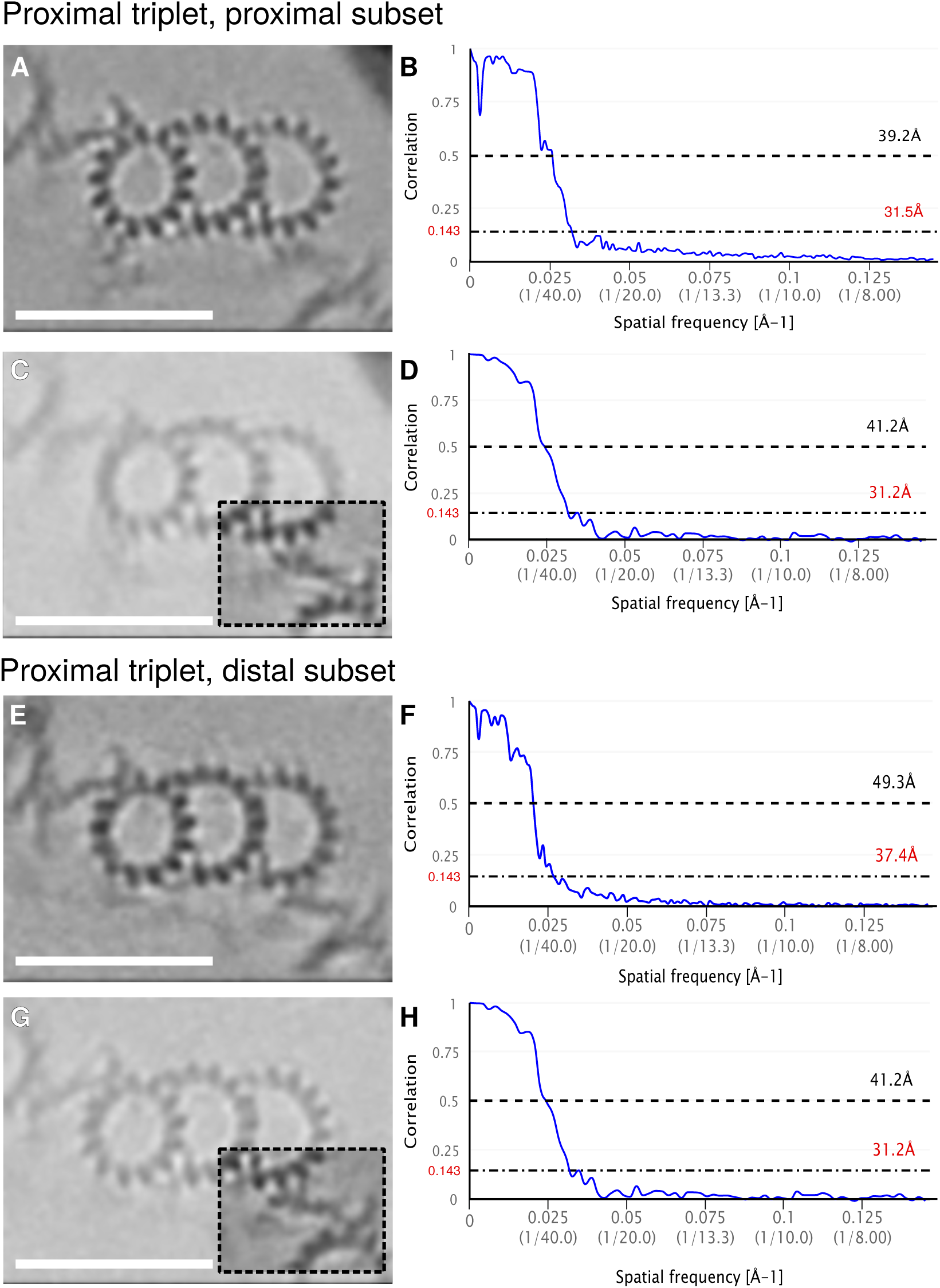
Resolutions of the subtomogram averages generated from the *P. tetraurelia* proximal centriole. (A, C, E, G) Z-projections of the obtained 3D maps (left) and their corresponding resolutions estimation by FSC curve (right, panel B, D, F, H). Initial maps were obtained by averaging the entire microtubule triplet from only the most proximal region (A) or the more distal part of the proximal region (E). Additional maps were made by local refinement of the A-C linker (dashed squared area) (C, G). Scale bars, 50 nm.

## Materials and Methods

### *Paramecium tetraurelia* centriole isolation and cryo-electron tomography

*P. tetraurelia* cortical units were isolated from two different strains, the wild type reference strain d4-2 and and Δ-CenBP1, as previously described (Le Guennec et al., 2020). Briefly, isolated *P. tetraurelia* centrioles were diluted with 1:1 colloidal gold in 10 mM K-PIPES buffer. Five microliters were deposited on 300 mesh lacey carbon grid and blotted from the backside before plunging in liquid ethane using a manual plunge freezing system. Tomograms were acquired with SerialEM software (Mastronarde, 2005) on a 300 kV FEI Titan Krios equipped with a Gatan K2 summit direct electron detector. The tilt-series were recorded from approximately −60° to +60° (bidirectional, 2° steps, separated at −0°), using an object pixel size of 3.45 Å, a defocus around −5 μm and a total dose of 70-120 electrons /Å^2^.

### Culture and *in situ* tomography of *Chlamydomonas reinhardtii* cells

The *in situ* of FIB-milling of *C. reinhardtii* centrioles was performed in the *mat3-4* strain, as previously described (Le Guennec et al., 2020). In brief, 4 μl of *C. reinhardtii* cells were deposited onto 200-mesh copper EM (R2/1, Quantifoil Micro Tools) and vitrified using a Vitrobot Mark 4 (FEI Thermo Fisher Scientific). Cryo-FIB sample preparation was performed as previously described (Schaffer et al., 2017, 2015). The FIB-milled EM grids were transferred into a 300-kV FEI Titan Krios transmission electron microscope, equipped with a post-column energy filter (Quantum, Gatan) and a direct detector camera (K2 Summit, Gatan). Tomogram were acquired using SerialEM software (Mastronarde, 2005), with tilt series between −60° and + 60° (bidirectional, 2° steps, separated at −0° or −20°) and a total dose around 100 electrons/Å^2^. A subset of tilt-series were acquired with a dose-symmetric scheme (Hagen et al., 2017). Individual tilts were recorded in movie mode at 12 frames/s, at an object pixel size of 3.42 Å and a defocus of −5 to −6 μm.

### *Naegleria gruberi* centriole isolation and cryo-electron tomography

Centriole isolation and tomogram acquisition were performed as previously described in (Le Guennec et al., 2020). Briefly, the *N. gruberi* NEG strain were differentiated into flagellates (Fulton, 1977), and centrioles were isolated using a sucrose gradient. Isolated centrioles were then deposited onto 200-mesh copper EM grids coated with holey carbon (R3.5/1, Quantifoil Micro Tools) and plunge-frozen in a liquid ethane/propane mixture. Tilt-series were recorded using SerialEM (Mastronarde, 2005) on a 300 kV FEI Titan Krios transmission electron microscope, equipped with a direct detector camera (K2 Summit, Gatan) and a post-column energy filter (Quantum, Gatan). Tilt-series were bidirectional (2° steps, separated at −0° or −20°), and individual images were recorded in movie mode at 10 frames/s, with an object pixel size of 4.21 Å and a defocus of −5 to −8 μm.

### Human centriole isolation and cryo-electron tomography

Human centrioles were isolated from the human lymphoblastic KE-37 cell line as previously described, (Gogendeau et al., 2015) with modification described in (Le Guennec et al., 2020). In brief, 5 μl of isolated centrioles diluted 1:2 with colloidal gold in 10 mM K-PIPES buffer were deposited on 300 mesh lacey carbon grids, blotted from the backside and quickly vitrified in liquid ethane using a manual plunge freezing. Tomogram acquisition was performed a 300 kV FEI Titan Krios equipped with a Gatan K2 summit direct electron detector. Bidirectional tilt series (2° steps, separated at −20°) were acquired with SerialEM (Mastronarde, 2005). Each tilt was recorded in movie mode at 12 frames/s with an object pixel size of 3.42 Å and a defocus of −4 to −6 μm.

### Subtomogram averaging of the A-C linker

From 11 tomograms of *P. tetraurelia* centrioles, 16 centrioles contained an intact proximal region. The positions of microtubules triplets were picked and interpolated every 8.5 nm as described in (Le Guennec et al., 2020) along the region displaying the A-C linker structure. Using Dynamo (Castaño-Díez et al., 2012), 1941 subtomograms of 320 × 320 × 320 voxels were extracted, encompassing the microtubule triplet with its associated A-C linkers. Initially, the microtubule triplets were roughly aligned on the *Trichonympha* reference (EMD-2330) (Guichard et al., 2013). To discriminate between subtomograms from the most proximal region and from the most distal region, a mask was created around the A-B inner junction where either the pinhead (a proximal marker) or the inner scaffold (a more distal marker) lies. A multireference alignment job was launched on this region, allowing us to classify our set into two classes: the “proximal-proximal” class (n = 1042) and the “proximal-distal” class (n = 899). For each set, the average was generated as a reference for the next step. Each set was then divided into two independent halves and aligned for a few iterations to produce two averages. The resolution was estimated using Fourier shell correlation (FSC) with a cut-off at 0.143. One of the averages was bandpass filtered at this resolution and the two half-sets were aligned on this filtered map to generate the final map.

The new aligned set was then split again into two halves, each half was locally aligned on the A-C linker region of the final map. After the two halves were aligned and the resolution computed, they were aligned on a common filtered map as previously performed for the global map.

The global map and the A-C linker map were combined together as described in (Le Guennec et al., 2020) to generate a volume displaying two adjacent microtubule triplets connected through the A-C linker. This map was then binned by a factor of 2 and combined with a rotated duplicate of itself to form a structure of the complete nine-fold proximal region, as described in (Le Guennec et al., 2020).

### Subtomogram averaging of the cartwheel

#### *P. tetraurelia* cartwheel

From 7 tomograms, 10 intact cartwheels were extracted as subtomograms with dimensions of 420 × 420 × 420 voxels. For each cartwheel, 9 duplicates were generated, each of them was rotated by a multiple of 40° to produce 9 different orientations of the original cartwheel. Each new volume was then shifted by −25, 0 or +25 nm to position a different unit of the cartwheel in the center of the volume. For each cartwheel, 27 subtomograms were generated (9 orientations x 3 units), resulting in 270 subtomograms in total from 10 cartwheels. To reduce the noise, the subtomograms were filtered using the nonlinear anisotropic diffusion command of Bsoft (Heymann et al., 2008).

An initial reference was generated by taking a cartwheel and its 8 differently oriented copies and averaging them together. The 270 subtomograms were aligned on this reference using SPIDER (Frank et al., 1996). After a few iterations, the average generated was used as a new reference on which the original, filtered but not aligned, subtomograms were aligned. From the 270 subtomograms, 38 failed to correctly align and thus were removed from the final set, resulting in 232 subtomograms used for the averaging. Nine-fold symmetry was then applied on the generated map to increase the contrast of the volume.

#### *C. reinhardtii* cartwheel

From five bin2 tomograms, 5 cartwheels were extracted as subtomograms with dimensions of 210 × 210 × 210 voxels. For each cartwheel, 9 duplicates were generated, each duplicate was rotated by a multiple of 40° to generate 9 different orientations of the original cartwheel. Each rotated volume was then shifted by −25, 0, or +25 nm to position different units of the cartwheel in the center of the volume. From five cartwheels, 9 × 3 = 27 subtomograms were generated resulting in 135 subtomograms in total. To improve the contrast, subtomograms were binned by a factor 2.

The 135 subtomograms were first aligned on the *P. tetraurelia* cartwheel map previously generated. Out of the 135 subtomograms, 86 were correctly aligned and used to produce an average map. This map was filtered by applying 3 iterations of Gaussian filter (with a sigma value of 2). The originally unaligned subtomograms were then aligned on this filtered average. 102 subtomograms were correctly aligned and kept to generate the average map. Nine-fold symmetry was then applied on the generated map to increase the contrast of the volume.

### Symmetrization

Top views of centrioles were generated using a z-projection of few slices from the cryo-tomogram and processed with the ImageJ plugin CentrioleJ for circularization and symmetrization (Guichard et al., 2013).

The symmetrization of the CID region was performed by generating a z-projection of a proximal part centered on the CID. From this image, 9 duplicates were generated by applying rotation from 0 to 360 degrees with a step of 40 degrees using Bsoft (Heymann et al., 2008). The 9 rotated images were then averaged together using SPIDER (Frank et al., 1996).

### Transmission electron microscopy of *Naegleria gruberi* serial section

*N. gruberi* NEG cells were differentiated from amoebae into flagellates as described in (Le Guennec et al., 2020), following a standard protocol (Fulton, 1977). Cells were fixed 50-80 minutes after the initiation of differentiation in order to observe both procentrioles and mature centrioles. The cells were pelleted and resuspended in 60 mM HEPES, 4 mM CaCl_2_, 2.5% glutaraldehyde, pH 7.2 and fixed for 120 min at room temp (replacing the fixative with fresh solution after 40 minutes). Cells were washed 2x 5 min in 60 mM HEPES, 4 mM CaCl_2_, pH 7.2 and osmicated using 1 % OsO_4_ in distilled water for 75 min at 4 °C. Cells were washed 3x 10 min in distilled water before en bloc staining in 1 % uranyl acetate in distilled water overnight at 4 °C. After washing 3x 10 min in distilled water, the cells were embedded in 1 % Agar noble (BD Difco, Sparks, MD, USA). Dehydration in ethanol, infiltration with Epon 812 (Serva Electrophoresis, Heidelberg, Germany) and final embedding was performed following standard procedures. Ultrathin serial sections (nominal 60 nm thickness) were cut with a diamond knife (type ultra 35°; Diatome, Biel, Switzerland) on an EM UC6 ultramicrotome (Leica, Wetzlar, Germany) and mounted on single-slot Pioloform-coated copper grids (Plano, Wetzlar, Germany). Sections were stained with uranyl acetate and lead citrate (Reynolds, 1963) and viewed with a JEM-2100 transmission electron microscope (JEOL, Tokyo, Japan) operated at 80 kV. Micrographs were acquired using a 4K charge-coupled device camera (UltraScan 4000; Gatan, Pleasanton, CA) and Gatan Digital Micrograph software (version 1.70.16.).

## Notes

### Competing Interest Statement

The authors have declared no competing interest.

